# scLinguist: A pre-trained hyena-based foundation model for cross-modality translation in single-cell multi-omics

**DOI:** 10.1101/2025.09.30.679123

**Authors:** Zhaoyu Fang, Ziyang Miao, Jianhui Lin, Yuying Xie, Jiliang Tang, Jiayuan Ding, Min Li

**Affiliations:** School of Computer Science and Engineering, Central South University, Changsha, China; Department of Computer Science and Engineering, Michigan State University, East Lansing, MI, USA; Department of Statistics & Probability, Michigan State University, East Lansing, MI, USA; Department of Computational Mathematics, Science and Engineering, Michigan State University, East Lansing, MI, USA

## Abstract

Single-cell multi-omics provides complementary insights into cellular states, heterogeneity, and regulatory programmes. However, paired assays remain costly, low-throughput, and technically challenging, whereas large-scale single-modality data such as scRNA-seq are abundant but do not capture protein-level biology. Here we present scLinguist, a foundation model for cross-modality translation introduces a three-stage framework: self-supervised pretraining on large-scale unimodal datasets to learn modality-specific representations, post-pretraining on limited paired data to capture cross-modality relationships, and inference to predict missing modalities (e.g., protein from RNA) in fine-tuning or zero-shot settings. Systematic benchmarking shows that scLinguist consistently outperforms state-of-the-art methods in predicting protein abundance from RNA across diverse biological contexts. It achieves high predictive performance while preserving cellular heterogeneity and further enables mechanistic and generalizable inference under simulated gene perturbations. Furthermore, scLinguist exhibits strong transferability across health states and datasets. By leveraging abundant unimodal data and minimizing dependence on paired assays, scLinguist establishes a scalable and versatile framework for cross-modality translation in single-cell analysis.

## Introduction

Single-cell sequencing has revolutionized our understanding of cellular systems, enabling high-resolution characterization of identity, dynamics, and heterogeneity. Technologies such as single-cell RNA sequencing (scRNA-seq) for transcriptomes, single-cell Assay for Transposase-Accessible Chromatin using sequencing (scATAC-seq) for chromatin accessibility, and single-cell proteomics for protein abundance provide complementary molecular perspectives that, together, offer a more comprehensive view of cellular regulation than any single modality alone (Baysoy et al. 2023). Yet cellular function is inherently multimodal, with chromatin accessibility, RNA expression, and protein abundance serving as key contributors to cell states and interactions. Multi-omics assays, such as RNA–protein co-profiling (CITE-seq (Stoeckius et al. 2017), AbSeq (Shahi et al. 2017)) and chromatin accessibility–RNA assays (Paired-seq (Zhu et al. 2019), SHARE-seq (Ma et al. 2020)) yield deeper biological insights but remain costly, technically demanding, and low-throughput, making large, high-quality paired datasets extremely rare. In contrast, single-modality datasets, particularly scRNA-seq, are abundant in both scale and coverage (CZI Cell Science Program et al. 2025), offering a rich resource for computational modeling and predictive approaches.

Despite the widespread availability of scRNA-seq, RNA levels alone are unreliable proxies for protein abundance due to post-transcriptional regulation, translational control, and protein degradation dynamics (Li and Xie 2011). Proteins, as the primary effectors of biological processes, play essential roles in signaling, metabolism, and structural organization (Specht et al. 2021; Angelo et al. 2014), and their accurate quantification is crucial for understanding cell fate determination, disease progression, and therapeutic responses (Schoof et al. 2021). However, single-cell proteomics faces substantial limitations: antibody-based methods detect only a limited, predefined set of proteins, while mass spectrometry can cover a broader range (Huffman et al. 2023), including intracellular proteins and post-translational modifications, but suffers from low sensitivity, high noise, and high cost. As a result, large-scale, unbiased protein quantification remains challenging (Zhu et al. 2019; Gatto et al. 2023), motivating the development of computational approaches to infer protein abundance from readily available scRNA-seq data. Several deep learning methods have been developed to predict protein abundance from RNA expression (Wu et al. 2021; Lakkis et al. 2022; Tang et al. 2023; Wen et al. 2022), including totalVI (Gayoso et al. 2021), scArches (Lotfollahi et al. 2022), and scButterfly (Cao et al. 2024). totalVI employs a probabilistic generative model to learn a shared latent space between RNA and protein, facilitating denoising and imputation. scArches extends this framework by incorporating transfer learning, allowing adaptation to new datasets while preserving biological variability. scButterfly utilizes an adversarial learning strategy to align RNA and protein representations across datasets. Although these methods have shown promising results, they are not designed as foundation models, often require extensive paired training data, rely on predefined latent spaces, and exhibit limited generalization across diverse datasets. In contrast, scTranslator (L. Liu et al. 2023) adopts a pretraining strategy on large-scale paired bulk multi-omics data before fine-tuning on single-cell paired samples. While this increases model capacity, bulk data lack single-cell resolution and are affected by population averaging, which limits their ability to capture rare cell states and fine-grained heterogeneity. Moreover, scTranslator does not fully leverage the vast amount of available single-modality data. Collectively, existing approaches fall short of delivering a scalable, generalizable, and interpretable solution for cross-modality translation.

Here, we present scLinguist, a pre-trained foundation model for cross-modality translation in single-cell multi-omics. scLinguist introduces a three-stage framework: (1) self-supervised pretraining on large-scale single-modality single-cell datasets to learn rich modality-specific representations, (2) post-pretraining on limited paired RNA-protein data to capture accurate cross-modality translation, and (3) inference from RNA profiles to predict protein abundance, either by fine-tuning on specific paired datasets or in a zero-shot setting. This design enables scLinguist to leverage abundant single-modality data while effectively learning modality relationships from limited paired datasets. Systematic evaluation demonstrates that scLinguist achieves more accurate protein abundance predictions, better preserves cellular heterogeneity, and effectively removes batch effects, even in few-shot and zero-shot scenarios. In addition, scLinguist can perform mechanistic or perturbation-like inference, enabling robust predictions under simulated gene perturbations. Overall, scLinguist provides a scalable and versatile platform for RNA-to-protein translation, enhancing the ability to generate and interpret single-cell proteomic landscapes from transcriptomic data.

## Results

### scLinguist introduces a large-scale data–driven framework for single-cell multi-omics translation

scLinguist introduces a general three-stage learning paradigm for cross-modal inference in single-cell multi-omics analysis, inspired by multilingual machine translation in Natural Language Processing (NLP). In NLP, translation models are typically pretrained on a large-scale monolingual corpora and subsequently post-trained on paired bilingual data (Lewis et al. 2019; Y. Liu et al. 2020), enabling robust representation learning and cross-lingual generalization. A similar challenge arises in single-cell multi-omics: large-scale unimodal datasets (e.g., RNA or protein profiles) are abundant, whereas paired multi-omics measurements are relatively scarce. To address this imbalance, scLinguist establishes a generalizable staged training strategy that applies across modalities. In Stage 1, we pretrained RNA- and protein-specific encoders and decoders independently on large unimodal datasets to learn robust unimodal representations. In Stage 2, paired RNA–protein data were used to further train both the RNA encoder and protein decoder, while introducing a cross-modal adapter module that connects the RNA encoder to the protein decoder, thereby enabling cross-modal alignment. In Stage 3, the pretrained model supports RNA-to-protein inference, which can be performed in a zero-shot manner directly from the pretrained model or further improved by fine-tuning on downstream datasets (Fig. 1a). While this paradigm can in principle generalize to other modality pairs, in this work we specifically focus on RNA-to-protein translation.

**Figure 1:**
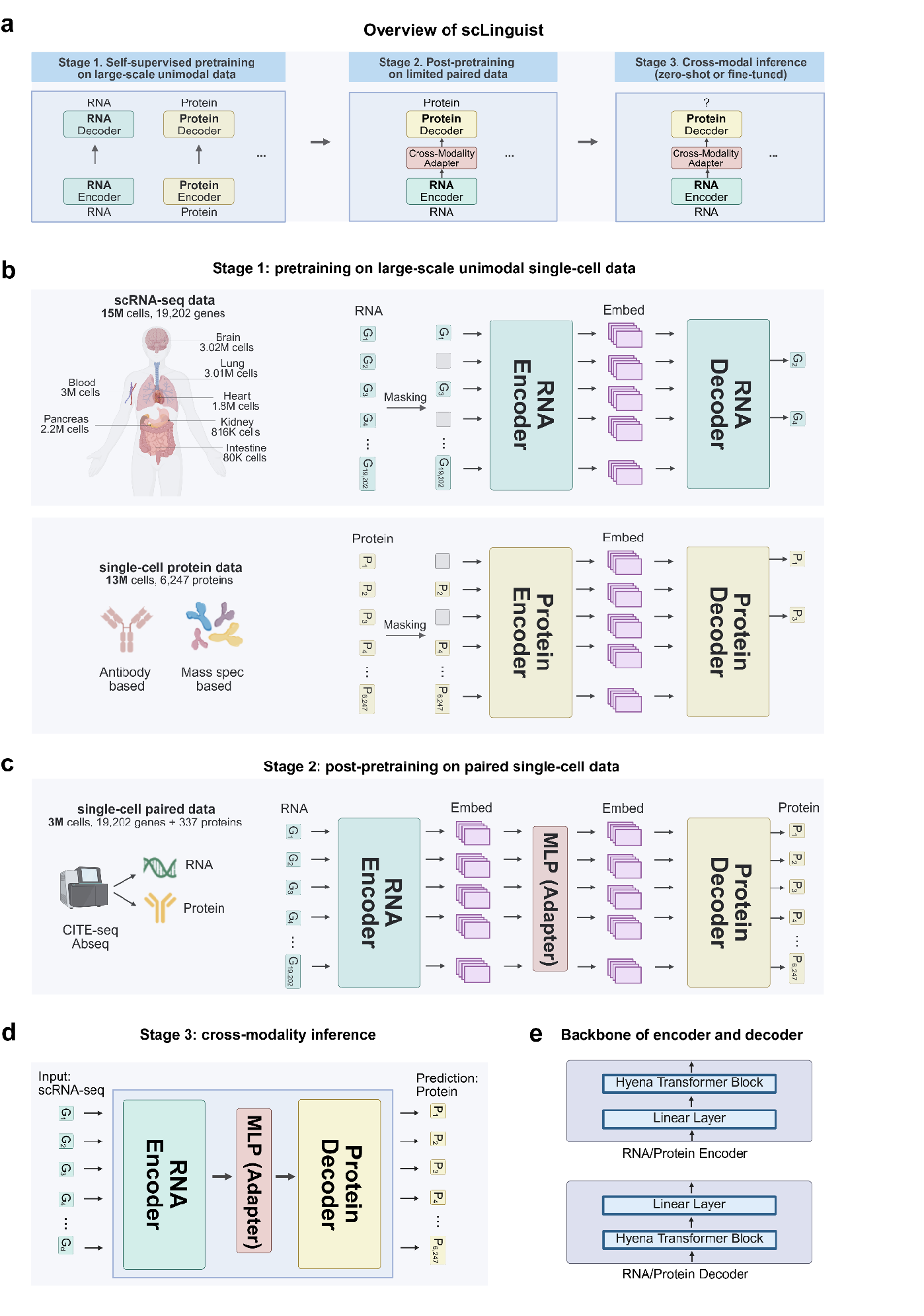
The schematic overview of scLinguist, a foundation model for single-cell multi-omics translation. a. **Overview**. scLinguist establishes a framework for predicting protein abundance from RNA profiles in single cells. The training proceeds in three stages. In Stage 1, RNA- and protein-specific encoders and decoders are pretrained in a self-supervised manner on large-scale unimodal datasets, enabling robust modality-specific representations. In Stage 2, the RNA encoder and protein decoder are further trained on paired RNA–protein single-cell datasets, with a cross-modal adapter connecting the two modalities. In Stage 3, the pretrained model performs RNA-to-protein inference, predicting protein abundance from RNA either in a zero-shot or fine-tuned setting. b. **Unimodality pretraining**. Each modality is trained as a denoising autoencoding task: a subset of input values is randomly masked, and the model reconstructs them by minimizing mean squared error (MSE). Both RNA and protein inputs are embedded as continuous-valued vectors to preserve quantitative expression information. c. **Cross-modality post-pretraining**. Paired RNA–protein data are used to connect a pretrained RNA encoder with a protein decoder through a multi-layer perceptron (MLP). The model predicts full protein expression profiles from RNA input by minimizing MSE between predicted and observed values. d. **Inference**. At inference time, scLinguist predicts the abundance of proteins from single-cell RNA input, exceeding the number of proteins typically measured with experimental technologies such as CITE-seq and AbSeq. e. **Backbone architecture**. All modality-specific encoders share the same architecture, and all modality-specific decoders share the same architecture. In the encoder, inputs are first projected through a linear embedding layer and then processed by stacked Hyena blocks. In contrast, the decoder first applies stacked Hyena blocks followed by a linear output layer.

To enable robust representation learning, we curated extensive pretraining datasets across both transcriptomic and proteomic modalities. Specifically, we assembled 15 million single-cell RNA sequencing (scRNA-seq) profiles from the CellxGene database (CZI Cell Science Program et al. 2025), covering a wide range of human tissues, including lung, brain, kidney, and blood. In parallel, we collected 13 million single-cell proteomic profiles from the SPDB database (Wang et al. 2024), incorporating both antibody-based and mass spectrometry-based technologies. For the cross-modal training stage, we further included 3 million paired RNA-protein profiles from SPDB, forming a high-quality corpus for learning cross-modality mappings. Together, these datasets together provide the scale and diversity required for training a generalizable foundation model for single-cell multi-omics.

scLinguist is built on a unified encoder-decoder architecture designed for both single-modality and cross-modality learning. The backbone of both the encoder and decoder is based on Hyena blocks (Poli et al. 2023) (Fig. 1e), which capture complex inter-feature dependencies, with a consistent architecture applied to both RNA and protein data. Unlike traditional approaches that discretize gene expression values into bins (Ding et al. 2025; Cui et al. 2024), scLinguist embeds continuous expression directly into high-dimensional latent vectors, preserving quantitative expression dynamics. During unimodal pretraining, the model is optimized as a denoising autoencoder (Fig. 1b). For RNA data, we implement a balanced dynamic masking strategy that includes both zero and nonzero values, encouraging the model to learn informative expression patterns rather than trivially reconstructing dropout. For protein data, we employ a simpler masking strategy that focuses on biologically meaningful reconstruction of protein-protein interactions. This phase enables the model to learn strong unimodal embeddings prior to cross-modal training.

In the post-pretraining phase, scLinguist performs cross-modal translation (e.g., RNA-to-protein) using a pretrained RNA encoder and protein decoder connected by an MLP-based cross-modal adapter. During this stage, both the encoder and decoder are further post-trained on paired multi-omics data, while the newly introduced adapter layer is jointly optimized to align the two modalities (Fig. 1c). Rather than relying on autoregressive decoding as in many sequence models, scLinguist performs single-step generation to predict the full protein expression profile in one forward pass. This non-autoregressive design improves scalability and inference speed, which is essential given the high dimensionality of omics data. The model is optimized using mean squared error (MSE) loss between predicted and observed protein values. In the inference phase (Stage 3), the pretrained model performs cross-modal prediction either in a zero-shot manner or with fine-tuning on downstream task datasets, enabling scLinguist to generalize beyond RNA-to-protein mapping to diverse modality pairs (Fig. 1d). At this stage, scLinguist can predict the expression of 6,247 proteins, far exceeding the capacity of widely used techniques such as CITE-seq (Stoeckius et al. 2017) or AbSeq (Shahi et al. 2017), which typically quantify fewer than 200 proteins.

Together, scLinguist provides a generalizable and data-efficient framework for multi-omics inference. Its non-autoregressive architecture ensures scalability and efficiency, making it suitable for large-scale applications in single-cell analysis. Leveraging this design, the framework can infer the expression of 6,247 proteins, far surpassing the capacity of conventional experimental assays. More broadly, scLinguist establishes a generalizable framework for multi-omics translation, trained on large-scale single-cell datasets and supporting robust inference across diverse omics layers.

### scLinguist consistently surpasses state-of-the-art methods in protein abundance prediction across varied learning settings

To comprehensively evaluate the performance of scLinguist in protein abundance prediction, we systematically compared it with four state-of-the-art (SOTA) models: totalVI (Gayoso et al. 2021), scArches (Lotfollahi et al. 2022), scButterfly (Cao et al. 2024), and scTranslator (L. Liu et al. 2023). Among these, totalVI and scArches were highlighted as the best-performing models in a recent comprehensive review (Hu et al. 2024). scTranslator is currently the only pretrained model specifically designed for protein prediction, while scButterfly represents a recently proposed cross-modal architecture. For fairness, we additionally standardized the input data for scButterfly to match the format used by scLinguist and scTranslator, referring to this variant as scButterfly_R. All models were benchmarked on four single-cell RNA-protein paired datasets (**Downstream task datasets**), following a comprehensive experimental design aimed at evaluating their robustness, generalizability, and adaptability.

We designed four experimental setups to evaluate generalization in realistic biological scenarios (**Fig. 2a**). Setting 1 employed a cell-type split, Setting 2 applied a random split, Setting 3 assessed few-shot learning with only 20 training cells, and Setting 4 evaluated zero-shot performance by directly applying the pretrained model to the test data. Under Setting 1, where models were tasked with predicting protein expression in unseen cell types during training, scLinguist consistently outperformed all SOTA methods across three key metrics: Pearson correlation coefficient (PCC), mean squared error (MSE), and maximum mean discrepancy (MMD) (**Fig. 2b**). On the BM dataset (Stuart et al. 2019), scTranslator achieved the second-best MSE and MMD, while scButterfly ranked second in PCC. On the BMMC dataset (Cao et al. 2024), scButterfly_R ranked second across all three metrics. Compared to the strongest comparison methods, scLinguist reduced MSE by an average of 33.28%, improved PCC by 8.82%, and reduced MMD by 26.73%, demonstrating superior accuracy, consistency with ground truth distributions, and ability to capture underlying expression trends. Visualization results further supported these findings. scLinguist accurately recovered biologically relevant protein signals such as CD45RA in memory B cells and CD56 bright NK cells, CD16 in CD16 monocytes, CD56 in NK cells, and CD38 in plasmablasts. In contrast, comparison methods often failed to capture these critical expression patterns. For example, scButterfly_R missed CD45RA in memory B cells, while scTranslator underestimated CD38 in plasmablasts (**Fig. 2c, Supplementary Fig. S1**).

**Figure 2.**
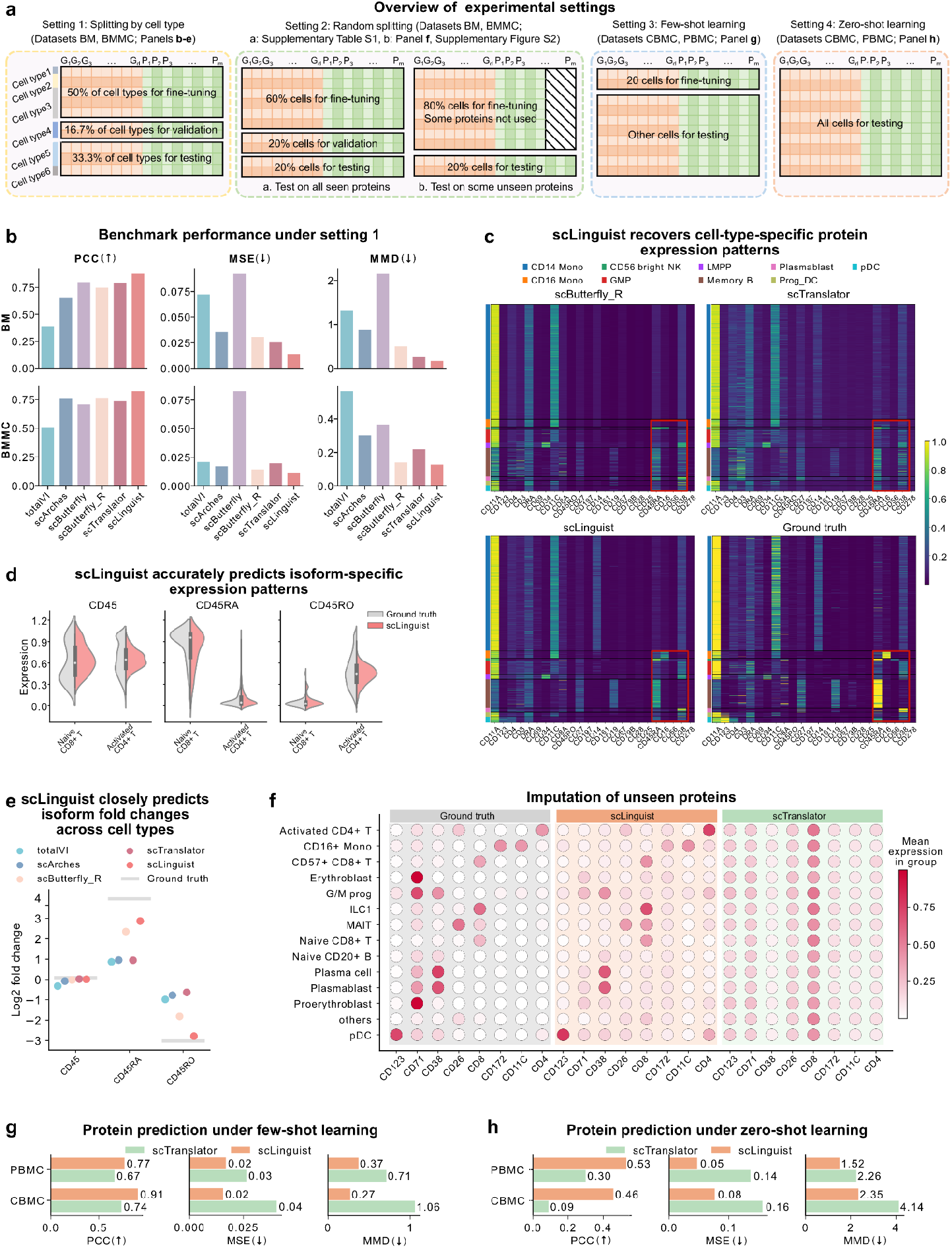
Benchmarking scLinguist across diverse protein prediction tasks. a. **Overview of experimental settings**. Four evaluation settings are designed to assess model generalizability: **Setting 1 (cell-type splitting):** 50% of cell types are used for training, 16.7% of unseen cell types for validation, and 33.3% of unseen cell types for testing. **Setting 2 (random splitting)**: (a) all cells are randomly shuffled and divided into 60% for training, 20% for validation, and 20% for testing; (b) 80% of cells are used for fine-tuning and the remaining 20% for testing, with the test set including proteins not present in training (unseen proteins). **Setting 3 (few-shot learning)**: only 20 cells are used for fine-tuning, while the remaining cells are used for testing. **Setting 4 (zero-shot learning)**: no fine-tuning is performed, and all data are used for testing. b. **Protein prediction benchmark performance under Setting 1, with test cell types entirely held out from training**. Performance of scLinguist, scTranslator, and scButterfly_R are evaluated on protein prediction from scRNA-seq across BM (Stuart et al. 2019) and BMMC (Cao et al. 2024) datasets using Pearson correlation coefficient (PCC), mean squared error (MSE), and maximum mean discrepancy (MMD) under the cell-type splitting scheme. c. **scLinguist recovers cell-type-specific protein expression patterns**. Heatmaps show predicted versus ground truth protein expression across major immune cell types, with color intensity indicating the density of protein expression values. d. **scLinguist accurately predicts isoform-specific protein expression patterns**. Violin plots comparing predicted and ground truth distributions of isoform-resolved proteins (CD45, CD45RA, CD45RO) across immune cell types. scLinguist accurately captures isoform-specific expression patterns, closely matching the ground truth profiles. e. **Recovery of isoform fold-change across cell types**. Log fold change (logFC) of CD45 isoforms between naïve CD8^+^ and activated CD4^+^ T cells, comparing predictions with true values. scLinguist closely tracks true differential expression patterns, outperforming baseline models in capturing isoform-specific variations. f. **Imputation results for eight proteins excluded from training in Setting 2b**. scLinguist accurately infers their cell-type-specific expression (e.g., CD123 in plasmacytoid dendritic cells (pDC), CD8 in ILC1 and CD8^+^ T cells), outperforming scTranslator. g. **Few-shot learning performance in Setting 3, where models are trained on only 20 paired cells**. scLinguist consistently outperforms scTranslator on PBMC (Peterson et al. 2017) and CBMC (Stoeckius et al. 2017) datasets. h. **Zero-shot learning performance in Setting 4, where pretrained models are directly applied to test data without any fine-tuning**. scLinguist consistently outperforms scTranslator on PBMC (Peterson et al. 2017) and CBMC (Stoeckius et al. 2017) datasets.

Single-cell proteomic data often include protein isoforms generated through alternative splicing, which, despite functional similarities, can exhibit markedly different expression across cell types (Su et al. 2023). The BMMC (Cao et al. 2024) dataset contains three CD45 isoforms—CD45, CD45RA, and CD45RO—offering an ideal case to test isoform-level prediction. CD45 is broadly expressed in both naïve CD8+ and activated CD4+ T cells, CD45RA is enriched in naïve CD8+ T cells, and CD45RO is specifically expressed in activated CD4+ T cells. scLinguist successfully captured these cell-type-specific patterns (**Fig. 2d**). To quantitatively validate this capability, we computed the log fold change (logFC) of predicted expressions between naïve CD8+ and activated CD4+ T cells and compared them with the true values. scLinguist’s predictions aligned most closely with the ground truth, outperforming all baselines in accuracy of isoform-specific predictions (**Fig. 2e**).

In Setting 2a, which involved random splitting of cells into training, validation, and test sets, scLinguist maintained its leading performance across all metrics on both BM (Stuart et al. 2019) and BMMC (Cao et al. 2024) datasets (**Supplementary Table S1**). Compared to the next-best models, scLinguist achieved an average reduction of 11.87% in MSE, a 1.05% increase in PCC, and a 4.97% reduction in MMD. The model’s consistent performance under both challenging (Setting 1) and standard (Setting 2a) conditions underscores the robustness of its pretraining framework and its strong generalization capability. To simulate real-world challenges where some protein targets are completely unobserved during training, we designed Setting 2b by excluding eight specific proteins from the training process and directly evaluating the model’s imputation performance on BMMC dataset (Cao et al. 2024). Since most models are incapable of predicting unseen proteins, this evaluation was limited to the pretrained scLinguist and scTranslator. scLinguist again outperformed scTranslator across all evaluation metrics (**Supplementary Fig. S2**). Heatmaps visualizing the imputed proteins revealed that scLinguist accurately captures the cell-type-specific protein expression patterns. For example, accurately predicting high CD123 in plasmacytoid dendritic cells (pDC) and CD8 in ILC1 and CD8+ T cells. (**Fig. 2f**). In contrast, scTranslator failed to clearly separate protein expression across relevant cell types, underscoring scLinguist’s superior ability to impute missing modalities.

To assess the model’s adaptability under limited supervision, we conducted both few-shot and zero-shot learning experiments. In Setting 3, we evaluated the few-shot learning capability of the models by training them on only 20 paired cells. scLinguist consistently outperformed scTranslator on both the PBMC (Peterson et al. 2017) and CBMC (Stoeckius et al. 2017) datasets (**Fig. 2g**). In the more challenging zero-shot scenario Setting 4, we directly applied the pretrained model to unseen test data without any downstream fine-tuning. Despite the absence of task-specific training, scLinguist successfully captured meaningful expression patterns and outperformed scTranslator across evaluation metrics (**Fig. 2h**). Moreover, scaling up the pretraining data consistently improved scLinguist’s zero-shot performance (**Supplementary Fig. S3**). These results highlight the strength of scLinguist’s pretraining framework, which enables it to generalize well and make accurate predictions even with minimal or no paired data.

In summary, scLinguist consistently outperforms existing SOTA methods across all learning settings. Its pretrained architecture enables robust generalization to unseen cell types, accurate imputation of unobserved protein targets, and strong performance under both few-shot and zero-shot learning conditions. These capabilities make scLinguist a powerful and scalable tool for real-world single-cell multi-omics applications, particularly in biologically complex or data-limited scenarios.

### scLinguist enables accurate protein prediction while preserving cellular heterogeneity

To evaluate how well predicted protein expression preserves biologically meaningful cellular heterogeneity, we systematically assessed the translation performance of different models through a series of downstream analyses, including batch correction, and cell type classification (**Fig. 3a**). Given that the BM (Stuart et al. 2019) and BMMC (Cao et al. 2024) datasets exhibit batch effects, we also examined each model’s ability to correct for these while preserving meaningful cellular structure.

**Figure 3.**
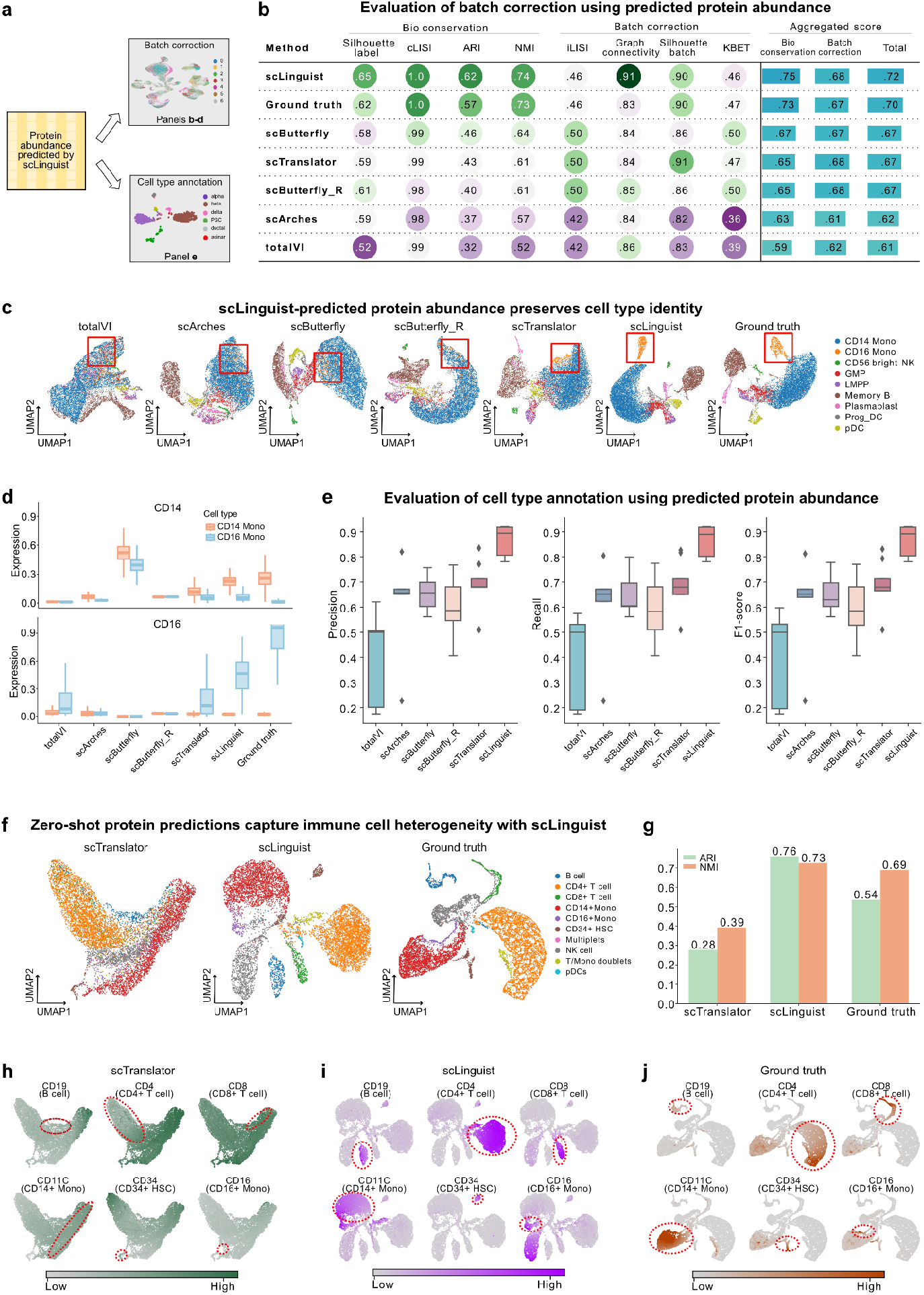
scLinguist enables accurate protein prediction while preserving cellular heterogeneity. a. **Overview of downstream analyses performed on predicted protein profiles, including batch correction and cell type classification**. b. **Quantitative evaluation of biological signal preservation and batch effect correction using predicted protein profiles, comparing scLinguist with other state-of-the-art methods**. Evaluations are conducted on BM (Stuart et al. 2019) and BMMC (Cao et al. 2024) datasets. scLinguist consistently outperforms all comparison methods across conservation and correction metrics. c. **UMAP visualization of predicted protein expression from the BM** (Stuart et al. 2019) **dataset (first test fold), colored by true cell types**. Only scLinguist and the ground truth clearly distinguish CD14^+^ and CD16^+^ monocytes (highlighted in red). d. **Average expression levels of CD14 and CD16 in CD14**^**+**^ **vs. CD16**^**+**^ **monocytes, comparing model predictions with ground truth protein data**. scLinguist recapitulates the expected differential expression patterns more accurately than other models. e. **Cell type classification based on predicted protein data**. scLinguist achieves the highest Precision, Recall, and F1 Score across comparison models. f. **UMAP visualization of zero-shot protein prediction, colored by true cell types**. scLinguist preserves clear cell type boundaries without fine-tuning, whereas scTranslator fails to separate key immune populations. g. **Clustering metrics under zero-shot predictions on the BMMC dataset** (Cao et al. 2024). scLinguist surpasses the performance of scTranslator and even exceeds ground truth. h. **Predicted protein expression by scTranslator, visualized by canonical marker expression**. scTranslator shows reduced specificity, with CD4 under-expressed in CD4^+^ T cells and diffusely expressed across other cell types, and CD8 expression lacking cell-type specificity. i. **Same as panel h, but for predictions from scLinguist**. scLinguist accurately recovers marker expression patterns, including CD19 (B cells), CD4 (CD4+ T cells) and CD8 (CD8+ T cells), CD11C (CD14+ monocytes), CD16 (CD16+ monocytes), and CD34 (hematopoietic stem cells). Co-expression of CD4 and CD11C in doublets is also correctly captured. j. **Same as panel h, but for the ground truth protein expression**. Ground truth confirms the expected expression of canonical surface markers, serving as a reference for evaluating model predictions.

We first evaluated how effectively predicted protein profiles from scLinguist, compared with state-of-the-art methods, preserved biological heterogeneity and mitigated batch effects across BM (Stuart et al. 2019) and BMMC (Cao et al. 2024) datasets (**Fig. 3b**). scLinguist achieved the highest overall performance, with strong scores across biological conservation metrics (Silhouette, cLISI, ARI, NMI) and batch correction metrics (iLISI, graph connectivity, silhouette, kBET), yielding the best aggregated score (0.72) and even surpassing the ground truth (0.70). By contrast, scButterfly and scTranslator showed moderate performance, while scArches and totalVI performed poorly, reflecting weaker correction and reduced biological fidelity. These results highlight scLinguist’s ability to jointly preserve cellular identity and mitigate batch effects more effectively than existing approaches.

We further used the BM (Stuart et al. 2019) dataset to show scLinguist’s superior ability to preserve cellular heterogeneity. On the first test fold, UMAP visualization of the predicted protein expression showed that only scLinguist and the ground truth data could clearly distinguish CD14^+^ monocytes from CD16^+^ monocytes, as highlighted in the red box (**Fig. 3c and Supplementary Fig. S4**). Other methods failed to accurately identify these two biologically important cell types.

To probe this further, we examined the expression patterns of the key surface markers CD14 and CD16 between the monocyte subset. As expected, the ground truth data showed CD14 enrichment in CD14^+^ monocytes and CD16 enrichment in CD16^+^ monocytes (**Fig. 3d**). Among the predicted profiles, only scLinguist and scButterfly correctly predicted the high CD14 expression in CD14^+^ monocytes. However, scTranslator incorrectly predicted higher CD14 levels in CD16^+^ monocytes. For CD16, although scLinguist, scTranslator, and totalVI all predicted the correct trend, the latter two exhibited substantially smaller expression differences. This explains why only the protein expression predicted by scLinguist could successfully distinguish between these two monocyte subtypes.

We next benchmarked model performance on supervised cell type classification using predicted protein profiles. scLinguist achieved the best performance across all metrics— Precision, Recall, and F1 Score—with median values approaching 0.9 (**Fig. 3e**). In comparison, scArches, scButterfly, scButterfly_R, and scTranslator yielded moderate performance (median 0.6–0.7), while totalVI performed worst, with metrics near 0.5. These results highlight scLinguist’s ability to preserve discriminative features essential for accurate classification.

To evaluate robustness under zero-shot learning conditions, we analyzed the BMMC dataset without any downstream fine-tuning. UMAP embeddings based on scLinguist and scTranslator predicted protein profiles showed that only scLinguist maintained clear boundaries between major immune cell populations (**Fig. 3f and Supplementary Fig. S5**). In contrast, scTranslator failed to distinguish key cell types such as CD4 versus CD8 T cells and CD14^+^ versus CD16^+^ monocytes. Clustering metrics confirmed these findings, with scLinguist not only outperformed scTranslator, but even exceeded the ground truth protein data (**Fig. 3g**), further demonstrating its superior ability to preserve cellular heterogeneity.

Finally, we assessed biological relevance by visualizing canonical surface marker expression. scLinguist accurately predicted high expression of CD19 in B cells, CD4 in CD4^+^ T cells, CD8 in CD8^+^ T cells, CD11C in CD14^+^ monocytes, CD34 in CD34^+^ hematopoietic stem cells (HSC), and CD16 in CD16^+^ monocytes (**Fig. 3h–j**). Remarkably, in a doublets cell population annotated as expressing both T cell and monocyte markers, scLinguist accurately predicted simultaneous high expression of CD4 and CD11C in these cells, reflecting true biological complexity. In contrast, scTranslator exhibited significant errors: CD4 was under-expressed in CD4^+^ T cells and aberrantly high in CD14^+^ monocytes; CD8 was broadly expressed across most cell types, lacking specificity and failing to accurately reflect its biological distribution.

In summary, scLinguist consistently preserved fine-grained cellular heterogeneity in the predicted protein expression space. Through batch correction, and cell type annotation tasks, it demonstrated strong biological consistency under both standard and zero-shot settings.

### scLinguist enables mechanistic and generalizable inference under genetic perturbations

To evaluate whether scLinguist can effectively capture regulatory relationships between RNA and protein expression, we designed two types of perturbation experiments (**Fig. 4a**). The first is a pseudo-perturbation experiment, where the expression value of a target gene is manually set to zero in the scRNA-seq data, simulating a CRISPR/Cas9 (Pandey et al. 2025) knockout at the computational level. The model then predicts pre- and post-perturbation protein expression levels, and fold-change analysis is used to identify the most responsive proteins. This low-cost computational approach enables preliminary functional exploration in scenarios lacking wet-lab access (Lotfollahi, Wolf, and Theis 2019). The second is a real perturbation prediction experiment based on publicly available single-cell data involving gene knockouts and matched RNA-protein measurements (Papalexi et al. 2021). These two experiments assess both the model’s mechanistic understanding of gene-protein interactions and its generalization to biologically realistic perturbations.

**Figure 4.**
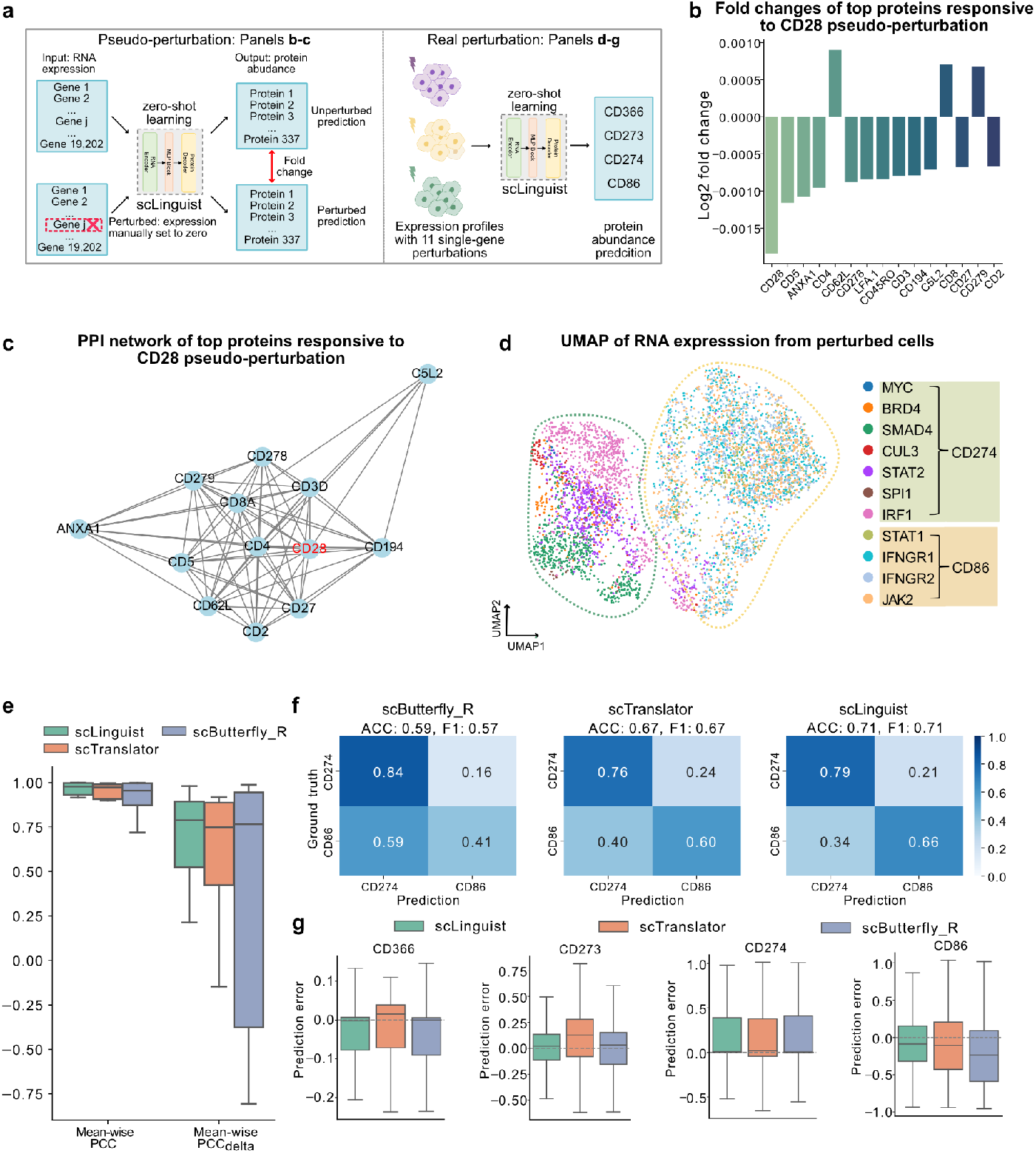
scLinguist enables mechanistic and generalizable inference under genetic perturbations. **a. Overview of two perturbation experiments: pseudo-perturbation (in silico gene knockout) and real perturbation (experimental gene knockouts)**. b. **Results of CD28 pseudo-perturbation**. The top 15 proteins with the largest predicted fold changes following in silico knockout of CD28 in PBMC (Peterson et al. 2017) dataset are shown. c. **Protein–protein interaction (PPI) network of the top 15 proteins responsive to CD28 pseudo-perturbation**. Edges represent known or predicted interactions from the STRING database; 12 out of 15 proteins are linked to CD28. d. **UMAP visualization of RNA expression profiles from the real perturbation dataset** (Papalexi et al. 2021). Cluster 1 (green box, right side): Enriched for perturbations that lead to high CD274 expression. Genes here include *MYC, BRD4, SMAD4, CUL3, STAT2, SPI1, IRF1*. Cluster 2 (orange box, left side): Enriched for perturbations that lead to high CD86 expression. Genes here include *STAT1, IFNGR1, IFNGR2, JAK2*. e. **Performance metrics in predicting perturbation-induced protein changes**. Two custom evaluation metrics are shown: mean-wise PCC, calculated as the Pearson correlation between predicted and true average protein expression across perturbation groups, and mean-wise PCC_delta_, calculated as the Pearson correlation between predicted and true changes in protein expression from the control to perturbed conditions. scLinguist achieves the highest accuracy across both metrics. f. **Confusion matrix for perturbation label classification based on predicted protein abundance**. A logistic regression classifier trained on predicted CD274 and CD86 levels distinguishes between CD274 and CD86 perturbation groups. scLinguist achieves the highest classification accuracy. g. **Boxplots of prediction errors for proteins across all perturbation conditions**. Prediction errors are calculated as the difference between true and predicted protein expression. scLinguist has the lowest median error across all proteins.

In the pseudo-perturbation experiments, we applied scLinguist to the PBMC (Peterson et al. 2017) dataset, setting the RNA expression of the immunoregulatory gene CD28 to zero. The model predicted changes in 337 protein expression levels and identified the 15 most responsive proteins (**Fig. 4b**). As expected, CD28 protein expression showed a significant decrease, confirming the model’s capacity to capture direct gene-protein relationships. Additionally, scLinguist predicted reduced expression of CD5, ANXA1, CD4, LFA-1, and CD3, all of which are associated with T cell activation and signaling (Beyersdorf, Kerkau, and Hünig 2015). In contrast, proteins such as CD62L, CD8, and PD-1 (CD279) showed increased expression, suggesting a cellular shift toward a quiescent or exhausted T cell state following CD28 inactivation. And the upregulation of PD-1 may reflect the induction of compensatory inhibitory mechanisms in response to impaired co-stimulatory signaling (Kim et al. 2021).

To evaluate the biological consistency of these predictions, we performed a STRING (Szklarczyk et al. 2019) network analysis, which revealed that 12 of the 15 responsive proteins have known or predicted interactions with CD28 (**Fig. 4c**). These results highlight scLinguist’s ability to infer meaningful mechanistic relationships within immunoregulatory networks. Additional pseudo-perturbation experiments targeting CD27, CD274 (PD-L1), CTLA4, and FOXP3 yielded similarly interpretable results (**Supplementary Fig. S6**).

To further evaluate predictive robustness, we used a real-world perturbation dataset (Papalexi et al. 2021) consisting of a non-perturbed control group and 11 single-gene perturbation groups (**Fig. 4d**). The dataset includes paired RNA expression and protein levels for CD366, CD274, CD273, and CD86. Dimensionality reduction and clustering based on RNA expression identified two major clusters: one associated with high CD274 expression (including MYC, BRD4, SMAD4, CUL3, STAT2, SPI1, IRF1 perturbations), and another with high CD86 expression (including STAT1, IFNGR1, IFNGR2, JAK2), consistent with classifications reported in the original study (Papalexi et al. 2021).

We trained scLinguist exclusively on the control data and applied it to predict protein expression in the perturbed groups. In addition to three standard evaluation metrics ( **Supplementary Fig. S7**), we introduced two additional metrics specifically designed to assess the model’s sensitivity to perturbation effects: mean-wise PCC, which measures the Pearson correlation between predicted and true average protein expression across different perturbation groups; and mean-wise PCC_delta_, which quantifies the correlation between predicted and true changes in protein expression from the control to the perturbed conditions (Cui et al. 2024). Across both metrics, scLinguist consistently outperformed comparison methods (**Fig. 4e**), demonstrating superior accuracy and best sensitivity to perturbation effects. In particular, both scLinguist and scTranslator showed clear advantages over scButterfly_R on the mean-wise PCC_delta_ metric, emphasizing their strength in modeling dynamic regulatory responses.

To evaluate the functional relevance of the predictions, we trained a logistic regression classifier using the predicted expression levels of CD274 and CD86 as features to classify into CD274 and CD86 perturbation labels. scLinguist achieved the highest classification accuracy, followed by scTranslator (**Fig. 4f**). While scButterfly_R performed comparably in the CD274 group, its performance declined notably in the CD86 group. Finally, we analyzed prediction errors for the four proteins and visualized them with boxplots (**Fig. 4g**). scLinguist exhibited the lowest prediction errors across all proteins, with median values close to zero. For CD274, both scLinguist and scButterfly_R delivered accurate predictions, whereas for CD86, scLinguist and scTranslator outperformed scButterfly_R, which is consistent with the classification results. These findings confirm scLinguist’s robustness, biological fidelity, and suitability for functional inference in perturbation-based single-cell analyses.

### scLinguist enables transfer across health states in scRNA-seq and tissue slides in spatial transcriptomics

To evaluate the generalization capability of scLinguist, we conducted experiments on both single-cell transcriptomics and spatial transcriptomics datasets. In the single-cell experiment, we used healthy human heart samples (Amrute et al. 2023) as the training set. For transfer evaluation, we randomly selected 20 cells from each of three types of heart failure patients (acutely infarcted, chronic ischemic, and non-ischemic cardiomyopathy) for few-shot fine-tuning, while the remaining heart failure cells were used for testing. In the spatial transcriptomics experiment, we selected two tissue slices from the same organ (tonsil), using one for training and the other for testing (**Fig. 5a**).

**Figure 5.**
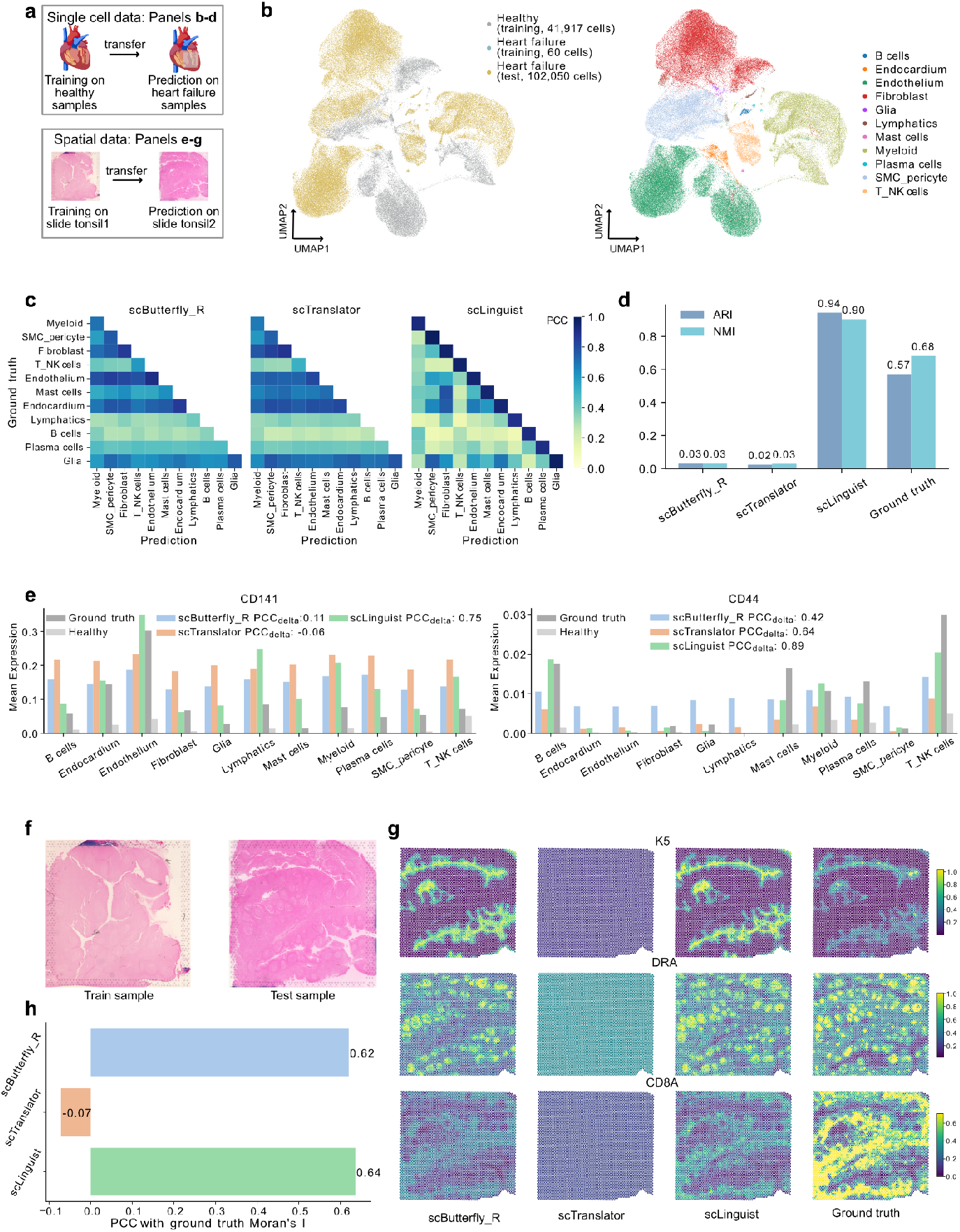
scLinguist enables transfer across health states in scRNA-seq and tissue slides in spatial transcriptomics. a. **Overview of the experimental setting**. In the scRNA-seq data setting, healthy human heart samples were used for training, and heart failure samples were used for few-shot fine-tuning and testing (Amrute et al. 2023). In the spatial transcriptomics experiments, two tonsil tissue sections were used, with one designated for training and the other for testing. b. **UMAP visualization of scLinguist-predicted protein expression in heart failure samples, integrated with true protein expression from healthy training samples and heart failure samples**. Cells of the same cell type cluster closely together, indicating strong consistency between predicted and true expression, as well as effective generalization to unseen patient data. c. **Heatmap of Pearson Correlation Coefficients (PCC) between predicted and true protein expression profiles across cell types**. For each predicted cell type, scLinguist achieves the highest correlation with its corresponding ground-truth type (diagonal dominance), outperforming other methods. d. **Clustering performance on predicted protein expression evaluated by Adjusted Rand Index (ARI) and Normalized Mutual Information (NMI)**. scLinguist shows superior preservation of cell-type heterogeneity compared to comparison methods. e. **Comparison of predicted average expression for five disease-associated proteins across cell types**. scLinguist successfully captures biologically consistent shifts in marker expression between healthy and heart failure conditions, while comparison methods fail to recover subtype-specific patterns. f. **H&E-stained tissue section used in spatial transcriptomics evaluation**. g. **Spatial distribution of predicted expression for marker proteins (K5, DRA, and CD8A)**. scLinguist successfully recapitulates known spatial localization patterns, including epithelial expression of K5, DRA enrichment at follicle borders, and CD8A localization in T cell–dense regions. h. **Correlation between predicted and ground-truth Moran’s I values**. scLinguist best preserves spatial heterogeneity, followed by scButterfly_R, with scTranslator failing to capture spatial organization.

We generated a UMAP visualization combining scLinguist predicted protein expression for heart failure samples with true expression from both healthy and heart failure training samples, which revealed clear separation of cell types and demonstrated strong generalization to previously unseen patient data (**Fig. 5b**). Quantitatively, scLinguist outperformed both scButterfly_R and scTranslator in MSE and PCC metrics (**Supplementary Fig. S8**). A heatmap of Pearson correlation coefficients (PCC) between predicted and true protein expression across cell types further confirmed that scLinguist consistently achieved the highest correlation with the corresponding ground truth cell type (**Fig. 5c**). For example, for the predicted Fibroblasts, the highest correlation was observed with the true Fibroblast profiles, followed by Mast cells and Endocardium, which is biologically consistent given their common origin in the connective tissue layer of the endocardium. In contrast, scTranslator and scButterfly_R show lower correlations in some cell types like B cells and Lymphatics. Moreover, their predictions for Endocardium and Endothelium exhibited high correlations with multiple unrelated cell types, indicating poor generalization capability.

To further evaluate whether the predicted protein expression preserves biologically relevant features after transfer, we performed clustering on the predicted protein profiles (**Supplementary Fig. S9**). scLinguist achieved the best performance in terms of Adjusted Rand Index (ARI) and Normalized Mutual Information (NMI), indicating superior preservation of cell-type heterogeneity (**Fig. 5d**). By comparison, scButterfly_R and scTranslator failed to clearly separate different cell types, further confirming their limited transfer ability.

We next evaluated whether scLinguist could capture biologically meaningful expression changes associated with disease. We selected five proteins known to differ significantly between healthy and heart failure samples, and compared the predicted average expression across cell types (**Fig. 5e, Supplementary Fig. S10**). For example, CD141 is highly expressed in T/NK cells in healthy samples but shifts to high expression in Endothelium and Endocardium in heart failure. scLinguist accurately captured this pattern. Similarly, CD44 is lowly expressed in healthy samples but upregulated in multiple immune-related cell types (B cells, T/NK, Mast, Plasma, and Myeloid) under heart failure conditions. scLinguist accurately predicted this trend, while scButterfly_R incorrectly predicted uniformly high expression of CD44 across all cell types, failing to capture subtype-specific patterns. We further computed the correlation (PCC_delta_) between predicted and true protein expression changes (difference from healthy samples) across cell types. scLinguist again achieved the highest PCC_delta_, confirming its robustness in cross-condition prediction.

To test the model’s generalizability beyond single-cell data, we evaluated scLinguist on spatial transcriptomics datasets. Although pre-trained solely on single-cell data, scLinguist was directly applied to predict protein expression using spatial transcriptomic inputs. Two tonsil samples were used, with one for fine-tuning and the other for testing (**Fig. 5f**). scLinguist outperformed other methods in MMD and MSE, while scButterfly_R performed slightly better in PCC (**Supplementary Fig. S11**).

We then visualized predicted protein distributions for several spatially informative markers, including K5, DRA, and CD8A. K5 is known to be highly expressed in epithelial regions, DRA in antigen-presenting cells at follicle borders, and CD8A in cytotoxic T cell-rich areas. scLinguist successfully captured their spatial enrichment patterns, closely matching ground truth (**Fig. 5g**). scButterfly_R produced comparable results, while scTranslator failed to resolve meaningful spatial organization, likely due to its reliance on bulk-level pre-training without cell-level resolution.

Finally, to quantify spatial coherence, we calculated Moran’s I (Chen 2013) for predicted protein expression and assessed its correlation with the ground truth Moran’s I values. scLinguist achieved the highest correlation (0.64), followed by scButterfly_R (0.62). scTranslator showed a negative correlation, confirming its failure to capture spatial heterogeneity (**Fig. 5h**).

In summary, scLinguist demonstrates strong generalization and transfer capability across both single-cell and spatial omics data. Even when trained solely on healthy samples, it is able to accurately predict protein expression in diseased tissues, supporting its applicability in cross-condition, cross-individual protein inference tasks.

## Discussion

Recent advances in single-cell technologies have enabled the generation of large-scale, multi-modal datasets capturing the molecular complexity of individual cells across diverse tissues and conditions. These data provide unprecedented opportunities for developing virtual models of cellular behavior, powered by generative AI. Inspired by the success of large language models (LLMs) in natural language processing, early single-cell foundation models such as Geneformer (Theodoris et al. 2023), scGPT (Cui et al. 2024), scFoundation (Hao et al. 2024), CellPLM (Wen et al. 2023) and Tabula (Ding et al. 2025) have adapted transformer-based architectures to biological contexts. However, these models primarily focus on transcriptomic data and do not fully leverage the rich cross-modal information available in multi-omics datasets, limiting their ability to model regulatory relationships across molecular layers.

In this work, we introduce scLinguist, a novel cross-modal foundation model based on an encoder–decoder architecture, designed to predict protein abundance from single-cell transcriptomic profiles. Drawing inspiration from multilingual translation models, scLinguist adopts a two-stage learning paradigm: it first performs modality-specific pretraining on large-scale unpaired omics data (e.g., RNA or protein) to capture intra-modality expression patterns, and then conducts post-pretraining on paired RNA–protein data to learn cross-modality mappings. This strategy enables the model to integrate knowledge from both data-rich and data-scarce scenarios, enhancing its generalizability and robustness across diverse biological contexts.

Built on a modular and flexible architecture, scLinguist specializes in cross-modal prediction, inferring one omics modality from another, such as predicting protein abundance from transcriptomic profiles. Its design allows straightforward extension to other modalities, including chromatin accessibility, DNA methylation, and spatial omics, enabling broad applicability in single-cell multi-omics studies. Comprehensive evaluations on benchmark datasets, including CITE-seq, REAP-seq, ECCITE-seq, and spatial data, demonstrate that scLinguist outperforms existing state-of-the-art methods in protein prediction and downstream tasks such as cell type annotation. Particularly in zero-shot and few-shot settings, the model exhibits strong generalization and transferability, effectively predicting biologically meaningful protein expression profiles even in limited-data scenarios.

As the scale and diversity of single-cell multi-omics datasets continue to expand, scLinguist offers a promising direction for constructing more comprehensive multi-omics virtual cells. Its ability to integrate across molecular layers and modalities provides a critical building block for generative frameworks that simulate cell states, perturbations, and responses. In the long term, such models may contribute to the realization of virtual laboratories and AI-driven biological discovery, transforming how we understand and manipulate cellular systems.

## Code availability

**scLinguist** (version: 1.0) is implemented with Pytorch Lightning and freely accessible at https://github.com/OmicsML/scLinguist.

## Methods

### Pretraining data collection and preprocessing

#### Data collection

For model pretraining, we assembled large-scale single-cell datasets spanning transcriptomic, proteomic, and paired RNA–protein measurements. The RNA data comprised 15 million cells sampled from the CELLxGENE database across diverse human tissues. Protein data (~13 million cells) were collected from the SPDB database, integrating antibody-based and mass cytometry-based technologies. In addition, ~3.6 million paired RNA–protein single cells from CITE-seq, Perturb CITE-seq, and ECCITE-seq were included to enable cross-modality training.

- **RNA data**. In this study, we utilized the 15 million-cell dataset provided by Ding et al. (Ding et al. 2025). This dataset originates from the CELLxGENE portal (https://cellxgene.cziscience.com) and is based on the version released on July 25, 2023. The CELLxGENE database comprises transcriptomic data from approximately 37.99 million human cells spanning various tissues. To ensure broad tissue representation, Ding et al. constructed a balanced subset by randomly selecting up to 3 million cells from each of the eight major tissue categories: intestine, pancreas, lung, heart, blood, kidney, brain, and a miscellaneous group labeled “others” (including tissues such as the spinal cord and spleen). For tissue types with fewer than 3 million available cells, all were included. The resulting 15 million-cell dataset preserves the biological diversity of the original collection while maintaining a balanced distribution across tissues, making it well-suited for robust model training.
- **Protein data**. We collected the single cell protein data from the SPDB database (https://scproteomicsdb.com). SPDB (Wang et al. 2024) integrates single-cell proteomics data generated via antibody-based technologies (e.g., CITE-seq, Abseq) and mass cytometry-based technologies (e.g., CyTOF). We downloaded the complete dataset from SPDB in November 2023. As CyTOF-derived data constituted the majority of the entries, we subsampled approximately 7.34 million cells from this modality to balance the distribution across different measurement technologies. Additionally, we included approximately 3.31 million cells from CITE-seq and 2.65 million cells from other technologies. The resulting dataset comprises approximately 13 million single cells and offers a more balanced representation across diverse experimental platforms.
- **Paired data**. Paired RNA-protein data were also obtained from the SPDB database and were used to facilitate cross-omics modeling and analysis. This dataset includes data generated using CITE-seq, Perturb CITE-seq, and ECCITE-seq technologies, comprising a total of approximately 3.6 million single cells.

#### Data preprocessing

For RNA data preprocessing, we first filtered out cells with fewer than 250 detected genes to exclude potential empty droplets and non-viable cells. After filtering, gene expression counts were normalized and log-transformed to reduce technical variations and enhance comparability across cells. We focused on human protein-coding genes, retaining a total of 19202 genes. If any genes were missing, their expression values were padded with zeros to ensure consistency across the dataset.

To address the substantial differences in protein abundance measurements between antibody-based and mass spectrometry-based single-cell proteomics platforms, we adopted a standardization approach inspired by scTranslator (L. Liu et al. 2023). Specifically, we applied Min-Max scaling to the count matrix on a per-cell basis, as formulated below:

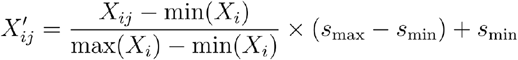

Where *i* represents the cell index, and the scaling factors were set to *s*_min_=10^−8^ and *s*_min_=1. This transformation preserves the dynamic range of protein expression while ensuring numerical stability for downstream analyses. We retained all unique proteins observed across the preprocessed datasets, totaling 6,427 proteins. To maintain consistency, missing values were zero-padded, as most datasets measured fewer than 100 proteins.

### scLinguist model overview

scLinguist is based on an encoder–decoder framework designed to leverage single-omics and paired multi-omics data. The pretraining process consists of three main stages. In Stage 1 (unimodal pretraining), the model is trained on individual RNA and protein datasets as a denoising autoencoder, analogous to a language translation model. Given a corrupted single-cell gene expression profile, scLinguist learns to reconstruct the original uncorrupted version by minimizing the mean squared error (MSE) between the predicted and true expression values, thereby capturing robust intra-modality representations. After Stage 1, the resulting modules, including the RNA encoder and decoder as well as the protein encoder and decoder, serve as pretrained components for the subsequent stage. In Stage 2 (cross-modal post-pretraining), the pretrained RNA encoder and Protein decoder are jointly fine-tuned on paired RNA–protein data to perform cross-modal translation. This stage enables the model to learn complex inter-modality relationships, effectively treating the task as a molecular translation problem. Finally, in Stage 3 (cross-modality inference), the model leverages the post-pretrained models to predict missing modalities under both fine-tuning and zero-shot settings.

#### Stage 1: Unimodal pretraining

In Stage 1, scLinguist performs unimodal pretraining by training two independent denoising autoencoders: an RNA autoencoder composed of an RNA-Encoder and an RNA-Decoder, and a protein autoencoder composed of a Protein-Encoder and a Protein-Decoder. Each autoencoder is trained exclusively on its respective modality to reconstruct masked input values, enabling the model to capture modality-specific intra-feature dependencies from RNA or protein expression profiles. While the overall network architecture, including the embedding layer, Hyena-based encoder–decoder backbone, and output projection, is kept identical across all encoders and decoders to maintain structural consistency, the parameters are not shared between RNA and protein modules. This design allows each module to learn modality-tailored representations while benefiting from a unified architectural framework. During pretraining, single-cell data are corrupted using a modality-specific masking strategy, and the model is optimized to minimize the mean squared error between the decoder’s output and the original uncorrupted data.

##### Architecture

This encoder–decoder architecture is used for both RNA and protein modalities, maintaining a consistent structural design while allowing independent parameters to capture modality-specific characteristics.

- **Encoder:** During the pretraining phase, the encoder takes either RNA or protein expression data as input. Although RNA and protein data differ in their original dimensionalities, for simplicity, we represent all modalities as a matrix *X* ∈ ℝ^*n* × *m*^, where *n* is the number of cells and *m* is the number of features, with each feature corresponding to a normalized expression value. Unlike natural language, where words are tokenized into discrete units and mapped to embeddings, gene expression values are continuous and cannot be directly tokenized. Previous studies have attempted to discretize expression levels into bins or use ranking-based methods (Bian et al. 2024), but these approaches risk losing important quantitative information in scRNA-seq data. To preserve the full quantitative content of gene and protein expression levels, we first expand the input tensor by introducing a singleton dimension:

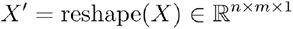

where each feature is treated as an individual token. Next, we apply a linear transformation to map the expression levels into a higher-dimensional embedding space:

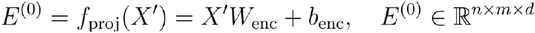

Where *W*_enc_ ∈ ℝ^1 × *d*^ is a learnable weight matrix that projects each feature individually into a *d*-dimensional space, and *b*_enc_ ∈ ℝ^*d*^ is a bias term. To efficiently model inter-feature dependencies, we employ a Hyena-based backbone (Poli et al. 2023). Hyena enables efficient long-sequence modeling and captures both local and global gene–gene relationships. Each feature column is treated as an individual input token, allowing the model to capture inter-feature correlations across genes or proteins. The input to the first Hyena layer is denoted as E^(0)^, and the output of the *l*-th Hyena block is computed as:

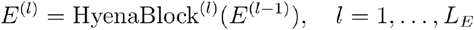

where L_E_ is the total number of Hyena blocks. Each Hyena block consists of a gated convolutional operator with long-range filters, followed by a feedforward layer. After passing through *L* stacked Hyena blocks, the final encoder representation is:

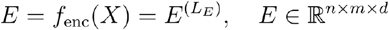

This encoder effectively transforms raw expression values into a structured representation, preserving the quantitative nature of the data while enabling the Hyena to learn intricate cross-feature dependencies.
- **Decoder**. The latent embeddings *E* are processed by *L*_*D*_ stacked Hyena blocks, refining the learned features:

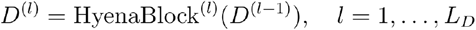

where HeynaBlock^(*l*)^ represents the *l*-th Hyena block in the decoder, and D^(0)^ = *E* is the output from the encoder. After passing through all Hyena blocks, the final transformed representation is denoted as:

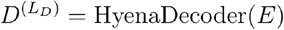

To map the learned embeddings back to the original expression space, we apply a linear projection:

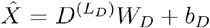

where *W*_*D*_ ∈ ℝ^*d* × 1^ is the weight matrix of the linear layer, *b*_*D*_ ∈ ℝ^*d*^ is the bias term, 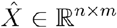 represents the reconstructed expression values. Thus, the overall decoder function can be expressed as:

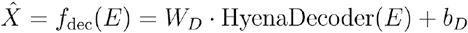

where *f*_dec_denotes the decoder function. The decoder can effectively refine the latent embeddings and reconstruct the original gene or protein expression values while preserving meaningful biological information.

##### Dynamic and balanced masking strategy

Single-cell RNA-seq data are highly sparse, with over 90% of expression entries being zeros. A naive random masking approach would predominantly mask zero-valued positions. Under such conditions, the model could trivially minimize reconstruction loss by predicting zeros for most masked entries, thus failing to learn meaningful biological representations. To overcome this limitation, we propose a dynamic and balanced masking strategy that, for each cell, masks 60% of the nonzero entries and an equal number of originally zero-valued entries. The total number of masked positions is dynamically determined based on each cell’s expression profile. This balanced masking forces the model to reconstruct both expressed and low-expressed genes, promoting robust embedding learning. Furthermore, it enables the model to distinguish true zeros, which reflect genuine low expression, from false zeros caused by dropout events. As a result, the model can more effectively capture biologically meaningful relationships and interactions between genes.

For protein data, which exhibit significantly lower sparsity compared to RNA, we adopt a simpler masking approach by randomly masking 40% of the measured protein expression values without balancing zeros. This simplified masking approach aligns with the dense nature of protein data and guides the model to focus on reconstructing biologically meaningful relationships and interactions between proteins.

##### Training objective

During single-modality pretraining, the random masking strategy is applied to the input expression matrix, where a portion of the expression values is masked (set to zero). The decoder is then trained to predict these masked values. The reconstruction loss is defined as the mean squared error (MSE) between the predicted expression values 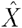 and the original, unmasked expression values *X*. Specifically, the loss function is given by:

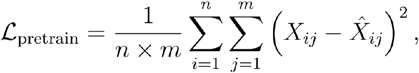

where *n* denotes the number of cells and *m* is the number of features. Minimizing ℒ _pretrain_ encourages the model to accurately recover the masked values, thereby promoting the learning of robust and informative latent representations.

#### Stage 2: Cross-modal post pretraining on paired data

Following the initial pretraining, we perform cross-modal post pretraining on paired RNA– protein datasets to capture the complex regulatory relationships between these two modalities. Inspired by advances in natural language processing, where leveraging pretrained language models has been shown to significantly enhance performance (Lewis et al. 2019; Y. Liu et al. 2020), we initialize our model with parameters obtained from pre-training on single-modality data: specifically, a pretrained RNA encoder and a pretrained protein decoder.

Our architecture adopts an encoder–cross modality adaptor–decoder framework, with RNA expression profiles as inputs and protein expression profiles as targets. To bridge the dimensional mismatch between RNA (dimension *g*) and protein (dimension *p*), we introduce a cross modality adapter implemented as a multi-layer perceptron (MLP) positioned between the encoder and decoder. This MLP projects and aligns the latent representations of RNA to the protein feature space, enabling effective cross-modal translation.

##### Architecture

Formally, given an RNA expression vector *X* ∈ ℝ^*n* × *g*^ (for *n* cells), the model processes it as follows:

- **RNA Encoder**: A pre-trained encoder *f*_*enc*_ extracts high-level RNA representations (as defined in Section Architecture):

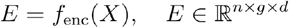

Here, *E* represents the encoded RNA features, where *d* denotes the dimension of the learned representation.
- **Cross-modality Adaptor**: A two-layer MLP transforms the RNA embeddings into the protein feature space. This process consists of two steps: The first layer maps the RNA representations to a hidden space using a ReLU activation function:

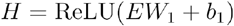

where *W*_1_ ∈ ℝ^*g* × *h*^ is the weight matrix, *b*_1_ ∈ ℝ^*h*^ is the bias vector, and *h* is the number of neurons in the hidden layer. The second layer projects the hidden representation *H* into the protein feature space:

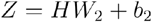

Where *W*_2_ ∈ ℝ^*h* × *p*^ is the weight matrix and *b*_2_ ∈ ℝ^*p*^ is the bias vector. The final output *Z* ∈ ℝ^*n*×*p*×*d*^ matches the protein feature dimensions, ensuring compatibility with the protein decoder. The overall transformation can be summarized as:

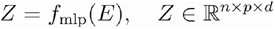
- **Protein Decoder**: A pre-trained decoder *f*_dec_ reconstructs protein expression levels from the projected representation *Z* (as defined in Section xxx):

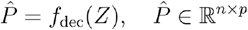

Where 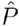 denotes the predicted protein expression matrix.

##### Training objective

During post pretraining, all model parameters are updated by minimizing the mean squared error (MSE) between the predicted and actual protein expression levels:

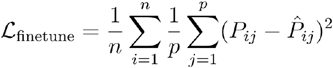

where P_ij_ and 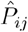 represent the ground-truth and predicted expression levels of protein *j* for sample *i*, while *p* is the number of proteins, and *n* is the number of cells.

By leveraging the representations learned during single-modality pre-training, the model effectively captures cross-modal relationships in paired RNA–protein data. The introduction of the Cross-modality Adaptor module ensures improved feature alignment, facilitating accurate translation between RNA and protein expression profiles.

#### Stage 3: Cross-modality inference

After completing unimodal pretraining (Stage 1) and cross-modal post pretraining on paired data (Stage 2), scLinguist proceeds to the final stage: cross-modality inference. In this stage, the model leverages the pretrained RNA encoder, cross-modality adaptor, and protein decoder to predict protein expression profiles from RNA-only datasets. This design enables the model to generalize beyond paired RNA–protein datasets, extending its applicability to large-scale scRNA-seq datasets where protein measurements are typically unavailable.

##### Architecture

During inference, the architecture follows the same encoder–adaptor–decoder pipeline as in Stage 2, but with only RNA input available. Given an input RNA expression profile, the pretrained RNA encoder transforms it into a latent representation:

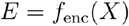

Next, the cross-modality adaptor projects the RNA embedding into the protein feature space:

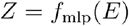

Finally, the protein decoder reconstructs the predicted protein expression values:

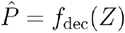

Here, 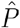 represents the predicted protein expression matrix for the input cells.

### Evaluation metrics

To quantitatively evaluate the generated cells, we employed three metrics: Pearson Correlation Coefficient (PCC), Maximum Mean Discrepancy (MMD), and Mean Squared Error (MSE). PCC was computed by flattening the expression matrices of both real and generated cells into vectors and calculating the correlation between them. For MMD, we applied principal component analysis (PCA) to both real and generated data when the number of proteins exceeded 100, using the resulting principal components for the discrepancy calculation; if the number of proteins was less than or equal to 100, the original expression data were used directly. MSE was calculated as the average of the squared differences between corresponding entries in the expression matrices of real and generated cells.

### Fine-tuning on downstream tasks and analysis

#### Batch correction

The batch correction performance of the predicted protein expression was evaluated using batch effect metrics from scib-metrics (Luecken et al. 2022). Specifically, we assessed biological conservation using Silhouette score on cell type labels (silhouette label), cell-wise local inverse Simpson’s index (cLISI), adjusted Rand index (ARI), and normalized mutual information (NMI). Batch correction effectiveness was quantified using inverse LISI (iLISI), graph connectivity, Silhouette score on batch labels (silhouette batch), and k-nearest neighbor batch effect test (kBET).

To enable clustering-based evaluation, Leiden (Traag, Waltman, and van Eck 2019) clustering was applied with the resolution parameter adjusted such that the number of resulting clusters matched the true number of cell types. We benchmarked our approach against totalVI, scArches, scButterfly, and scTranslator on BMMC (Cao et al. 2024) and BM (Stuart et al. 2019) datasets, providing a comprehensive assessment of batch correction performance and biological conservation.

#### Cell type annotation

To perform cell type annotation, we trained an MLP classifier on true protein expression data. This classifier takes protein expression profiles as input and outputs categorical predictions for cell types. The classification model is optimized using the cross-entropy loss, formulated as:

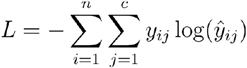

where *n* is the number of cells in a batch, *c* represents the total number of cell type classes, *y*_*ij*_ is the one-hot encoded ground truth label for the *i*-th cell belonging to class *j*, and ŷ _*ij*_ is the predicted probability of the *i*-th cell being assigned to class *j*.

After training, this classifier was used to evaluate the protein expression profiles predicted by different methods, including scLinguist, totalVI, scArches, scButterfly, and scTranslator, to generate cell type predictions. We benchmarked the classification performance on two datasets, BMMC (Cao et al. 2024) and BM (Stuart et al. 2019), using precision, recall, and F1 score as evaluation metrics, providing a comprehensive assessment of the predictive accuracy of each method.

#### Pseudo-perturbation

We employed the pretrained scLinguist model to investigate the impact of RNA-level gene pseudo-perturbation on protein expression. Specifically, we simulated gene knockout by setting the expression value of a target gene to zero in the input RNA matrix and calculated the log fold change in predicted protein abundance before and after the perturbation. In this study, we analyzed the effects of perturbing several immunoregulatory genes, including CD28, CD27, CD274 (PD-L1), CTLA4, and FOXP3. These pseudo-knockout experiments were conducted using the PBMC (Peterson et al. 2017) dataset, and protein-level responses were predicted for a panel of 337 proteins learned during post-pretraining. For each gene perturbation, we identified the top 15 proteins exhibiting the most significant changes in expression. To assess the biological relevance of the predictions, the top responsive proteins were further analyzed using the STRING database to identify known or predicted protein-protein interaction networks.

### Downstream task datasets

For downstream evaluation, we employed six publicly available single-cell multi-omics datasets and one spatial multi-omics dataset, all of which contain joint measurements of transcriptomic and proteomic data. To ensure consistency and comparability, we retained only those proteins that overlapped with the pretraining protein set, which may reduce the number of proteins compared to the original datasets.

#### BM dataset (GSE128639)

Based on CITE-seq technology, this dataset includes 30,672 human bone marrow-derived single cells with transcriptomic profiles (scRNA-seq) and measurements of 25 surface proteins, collected across two batches (Stuart et al. 2019). In this study, we retained 24 proteins that overlapped with the pretraining set.

#### BMMC dataset (GSE194122)

This CITE-seq dataset includes 90,261 bone marrow mononuclear cells (BMMCs) from 12 healthy donors across 12 batches, with measurements of 13,953 genes and 134 surface proteins. It also provides annotations for 45 immune cell subtypes (Cao et al. 2024). Originally used in the NeurIPS 2021 multimodal single-cell integration challenge, this dataset has been widely adopted in multimodal single-cell research (Luecken et al. 2021).

#### CBMC dataset (GSE10086)

Based on CITE-seq, this dataset contains 8,005 cord blood mononuclear cells (CBMCs) with measurements of 16,508 transcriptomic genes and 13 surface proteins (Stoeckius et al. 2017).

#### PBMC dataset (GSE100501)

Generated using REAP-seq, this dataset includes 4,330 peripheral blood mononuclear cells (PBMCs) with measurements of 21,005 transcriptomic genes and 44 surface proteins (Peterson et al. 2017). We retained 40 proteins that overlapped with the pretraining protein set.

#### Perturb dataset (GSE153056)

Generated using ECCITE-seq, this dataset integrates CRISPR perturbation screening, transcriptomic profiling, and surface protein detection at the single-cell level. It was designed to systematically explore the regulatory network of PD-L1 expression and includes various gene perturbation experiments (Papalexi et al. 2021).

#### Heart dataset (GSE218392)

Based on CITE-seq, this dataset includes 41,917 cells from healthy samples and 102,110 cells from three types of heart failure patients (acutely infarcted, chronic ischemic, and non-ischemic cardiomyopathy), with 257 proteins measured (Amrute et al. 2023).

#### Spatial dataset

Obtained from the Visium platform by 10x Genomics, this dataset includes two human tonsil samples. We used measurements of 27 proteins for analysis (10x 2023a, 2023b).

### Benchmarking methods

We benchmarked scLinguist against other state-of-the-art methods, including scTranslator, scButterfly, totalVI, and scArches. Additionally, we standardized the input data for scButterfly to match the format used by scLinguist and scTranslator, referring to this modified version as scButterfly_R.

#### scTranslator (L. Liu et al. 2023)

We used the official scTranslator implementation in PyTorch Lightning. We followed the authors’ standard preprocessing and training workflow (https://github.com/TencentAILabHealthcare/scTranslator) and obtained protein IDs using the provided mapping files. The model was initialized from a released 2M checkpoint and fine-tuned end-to-end on the training split for 6 epochs. Training used the package’s default optimizer, loss, and learning-rate scheduler; no other hyperparameters were changed.

#### scButterfly (Cao et al. 2024)

We employed the public scButterfly implementation in Python and followed the official tutorial (https://scbutterfly.readthedocs.io/en/latest/). Raw RNA and ADT count matrices were supplied as inputs; RNA was normalized and log-transformed with the package’s RNA_data_preprocessing, and ADT was processed with the authors’ e centered log-ratio (CLR) normalization after removing all-missing antibody channels to align panels. We then applied the default two-stage training recommended by the authors— autoencoder pretraining per modality followed by translator training with early stopping—using the default network depth and embedding size. Batch size and learning rate followed the library defaults (we used batch size 32 and learning rate 1e−3), and train/validation/test splits matched our benchmark setup.

#### scButterfly_R

It is a variant introduced to align the ADT preprocessing with that of scLinguist. This variant differs only in ADT preprocessing: instead of CLR, we applied feature-wise min– max scaling to [1e−8, 1] with explicit handling of missing values, keeping RNA preprocessing, architecture, optimizer settings, and the two-stage training schedule identical to scButterfly. Data splits and input format were held constant to isolate the effect of the ADT normalization scheme.

#### totalVI (Gayoso et al. 2021)

We used totalVI from scvi-tools (Gayoso et al. 2022) in Python, following the Multiome Benchmarking workflow (https://github.com/QuKunLab/MultiomeBenchmarking). To assess protein prediction, proteins in the held-out batch were masked during training and imputed at inference. The model was initialized with a normal latent distribution and a two-layer decoder and trained with the library’s default optimizer/early-stopping settings using the tutorial’s train/validation proportions. We then exported the latent representation (X_totalVI), protein foreground probabilities, and imputed protein means for the held-out batch. No other hyperparameters were changed.

#### scArches (Lotfollahi et al. 2022)

We used scArches in scvi-tools (Gayoso et al. 2022) with TOTALVI as the backbone, following the standard reference–query workflow. We used the authors’ recommended settings and trained with default optimization, using the train/validation proportions derived from sample counts. The saved reference was then used to load the query with freeze_expression=True for amortized transfer, and the query was fine-tuned for 200 epochs. We computed latent embeddings (X_totalVI) and imputed query-batch proteins via get_normalized_expression with transform_batch set to the reference batch (25 posterior samples, mean). No other hyperparameters were changed.

