## Supplementary materials for "scLinguist: A pre-trained hyena-based foundation model for cross-modality translation in single-cell multi-omics"

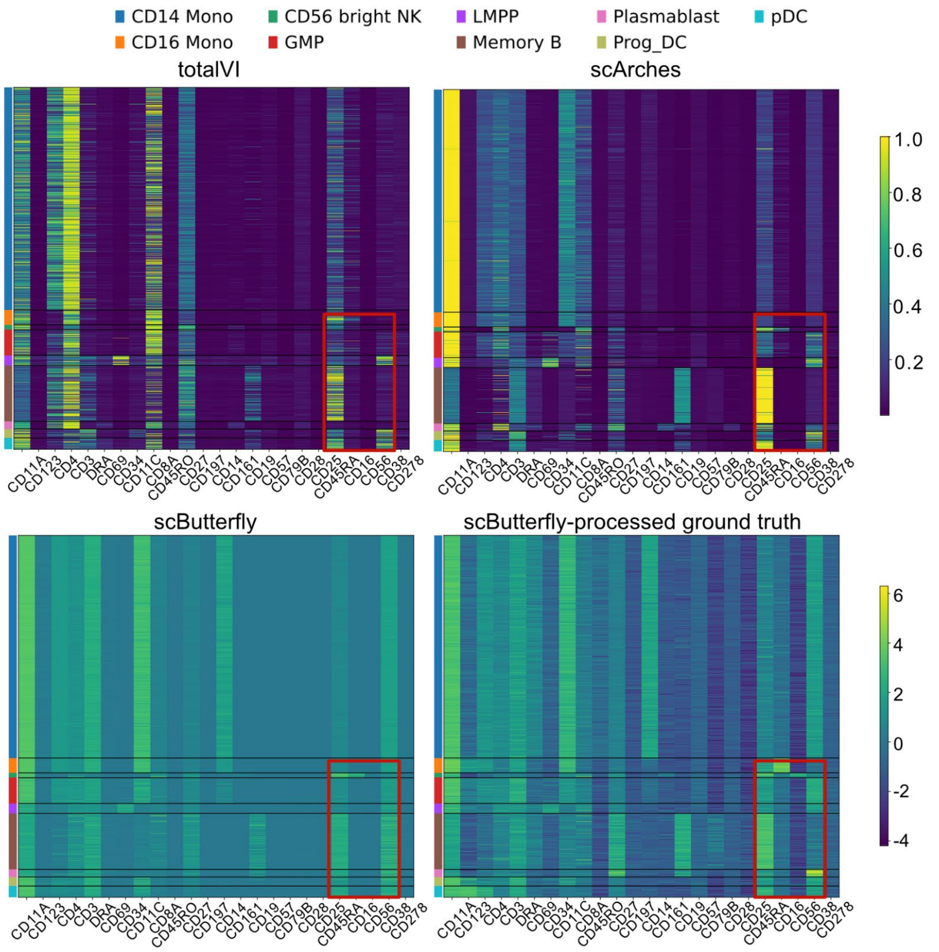


**Figure S1**. Heatmap displaying protein expression patterns across cell types predicted by totalVI, scArches, and scButterfly, alongside ground truth data processed with scButterfly.

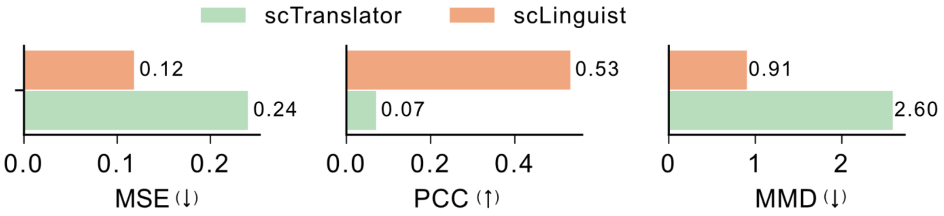


**Figure S2**. Quantitative comparison of model performance under Setting 2b (missing proteins) on the BMMC dataset.

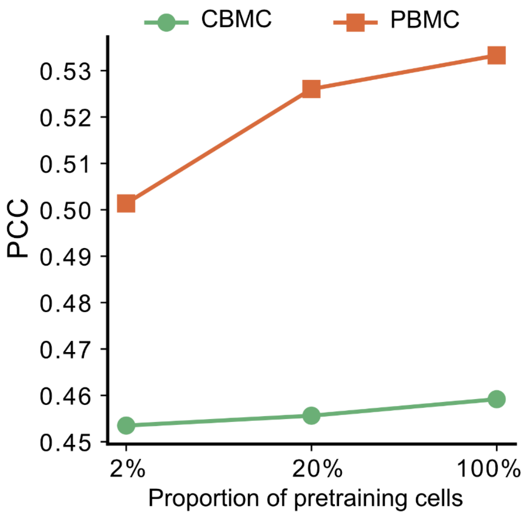


**Figure S3**. Scaling law analysis of scLinguist under zero-shot setting. scLinguist was first pretrained on 2%, 20%, or 100% of the available single-modality RNA and protein datasets, followed by post-pretraining on the same 300k paired RNA–protein cells. The resulting models were directly evaluated on PBMC (squares) and CBMC (circles) datasets without any downstream fine-tuning. The x-axis denotes the proportion of cells used during the initial pretraining, and the y-axis shows the prediction accuracy measured by Pearson correlation coefficient (PCC). The results show that scLinguist's pretraining strategy effectively enables zero-shot prediction, with performance improving as more pretraining data is used, highlighting the utility of the pretraining framework.

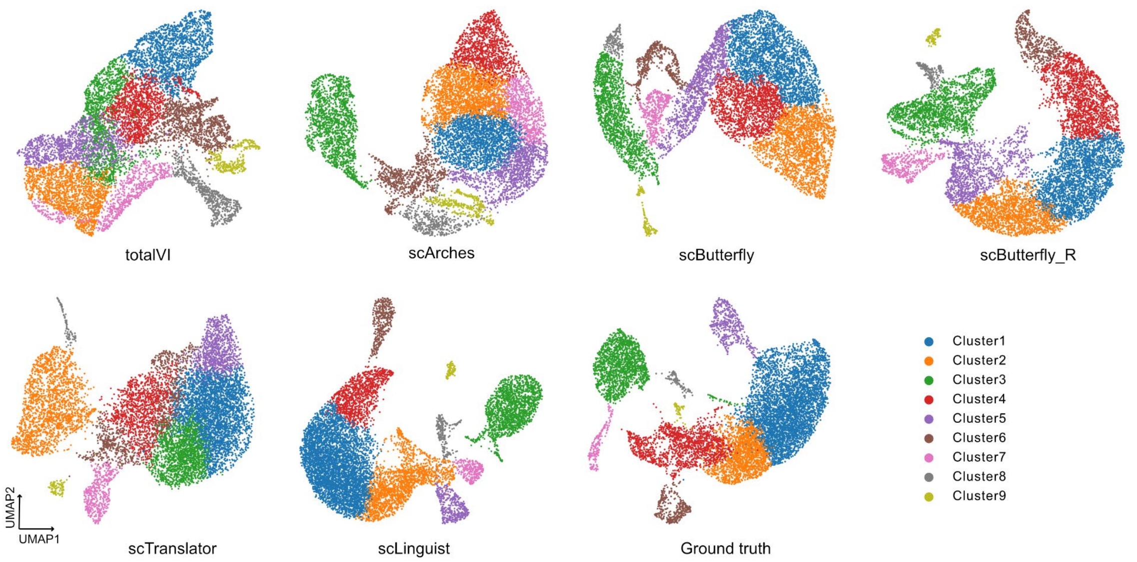


**Figure S4**. UMAP plots of predicted protein expression from different models along with ground truth data, colored by clustering labels.

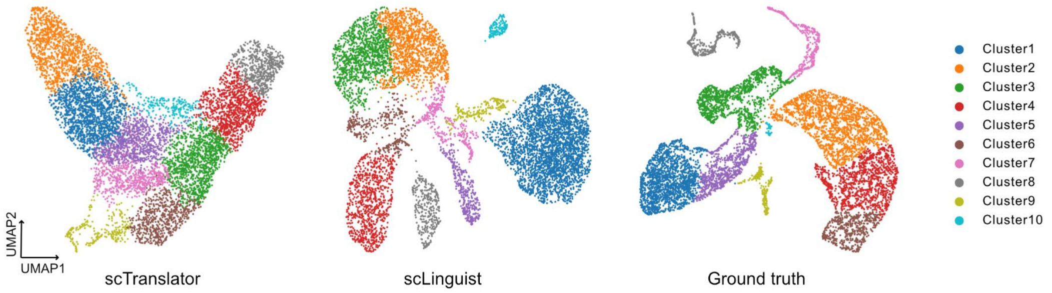


**Figure S5**. UMAP embeddings of protein expression predicted by scTranslator and scLinguist without fine-tuning, along with ground truth data, colored by clustering labels.

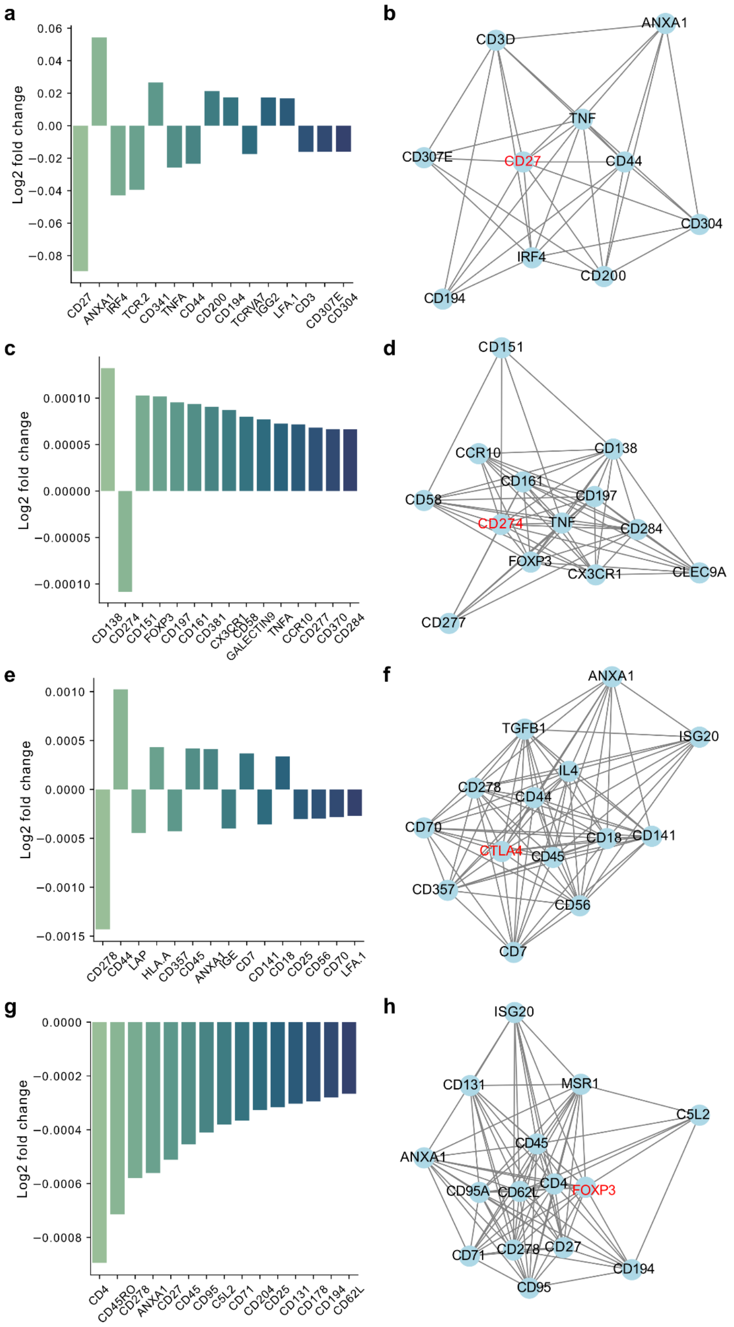


**Figure S6**. Predicted protein expression changes following in silico knockout of CD27, CD274 (PD-L1), CTLA4, and FOXP3. (a) Log fold changes of the top 15 proteins most affected by pseudo-perturbation of CD27 in the PBMC (Peterson et al. 2017) dataset. (b) STRING network analysis of the top 15 proteins responsive to CD27 perturbation, showing known interactions with CD27. (c) Top 15 proteins affected by CD274 pseudo-perturbation. (d) STRING network of CD274-responsive proteins. (e) Top 15 proteins affected by CTLA4 pseudo-perturbation. (f) STRING network of CTLA4-responsive proteins. (g) Top 15 proteins affected by FOXP3 pseudo-perturbation. (h) STRING network of FOXP3-responsive proteins.

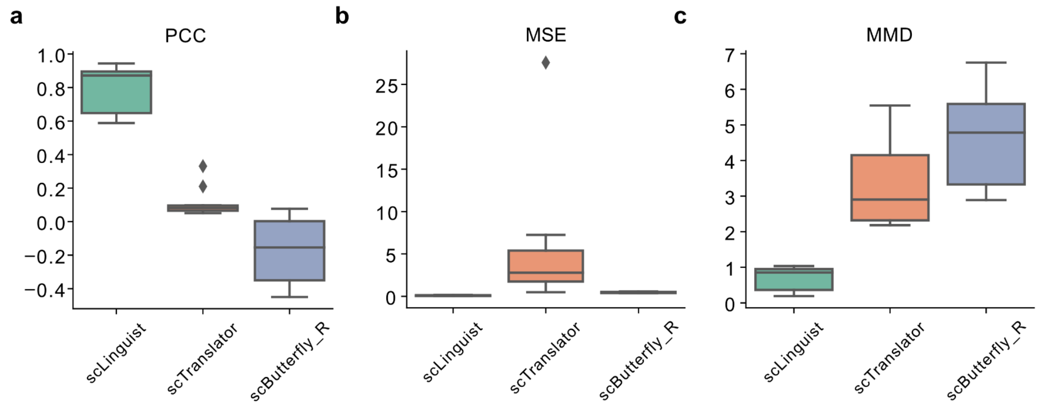


**Figure S7**. Evaluation metrics for model performance on real perturbation dataset (Papalexi et al. 2021).

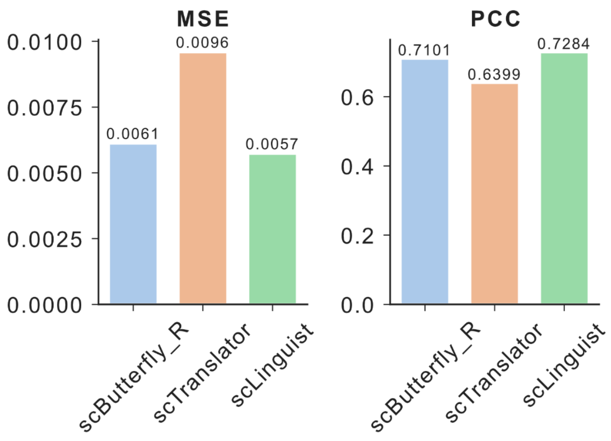


**Figure S8.** Evaluation metrics for protein prediction performance on heart failure single-cell data (Amrute et al. 2023). Bar plots comparing scLinguist, scButterfly_R, and scTranslator on Mean Squared Error (MSE) and Pearson Correlation Coefficient (PCC). scLinguist consistently achieves the best performance across both metrics, demonstrating superior predictive accuracy.

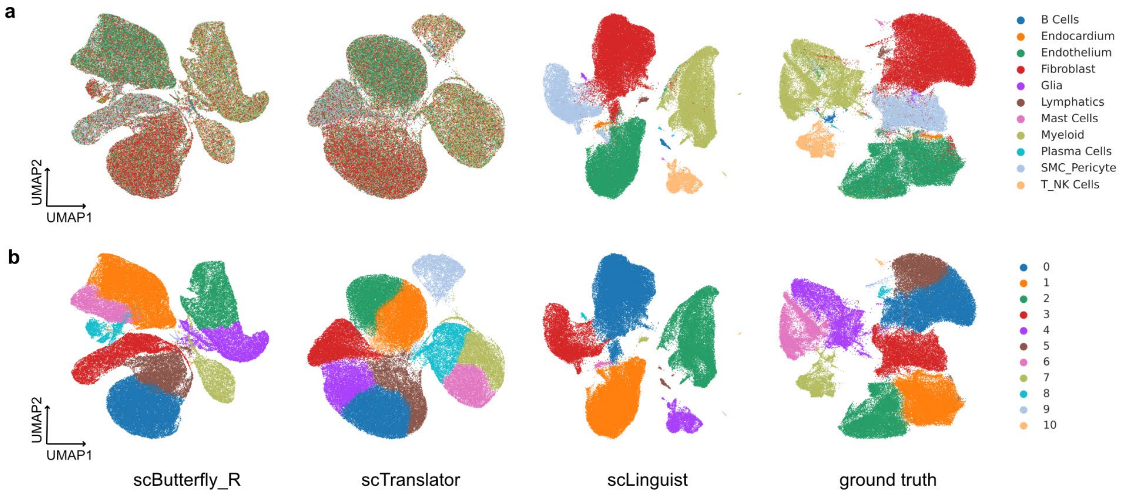


**Figure S9.** UMAP plots of protein profiles predicted by scLinguist, scButterfly_R, and scTranslator. (a) Cells are colored by ground-truth cell types. (b) Cells are colored by clustering results. scLinguist exhibits clear separation of cell types, suggesting strong preservation of biological heterogeneity.

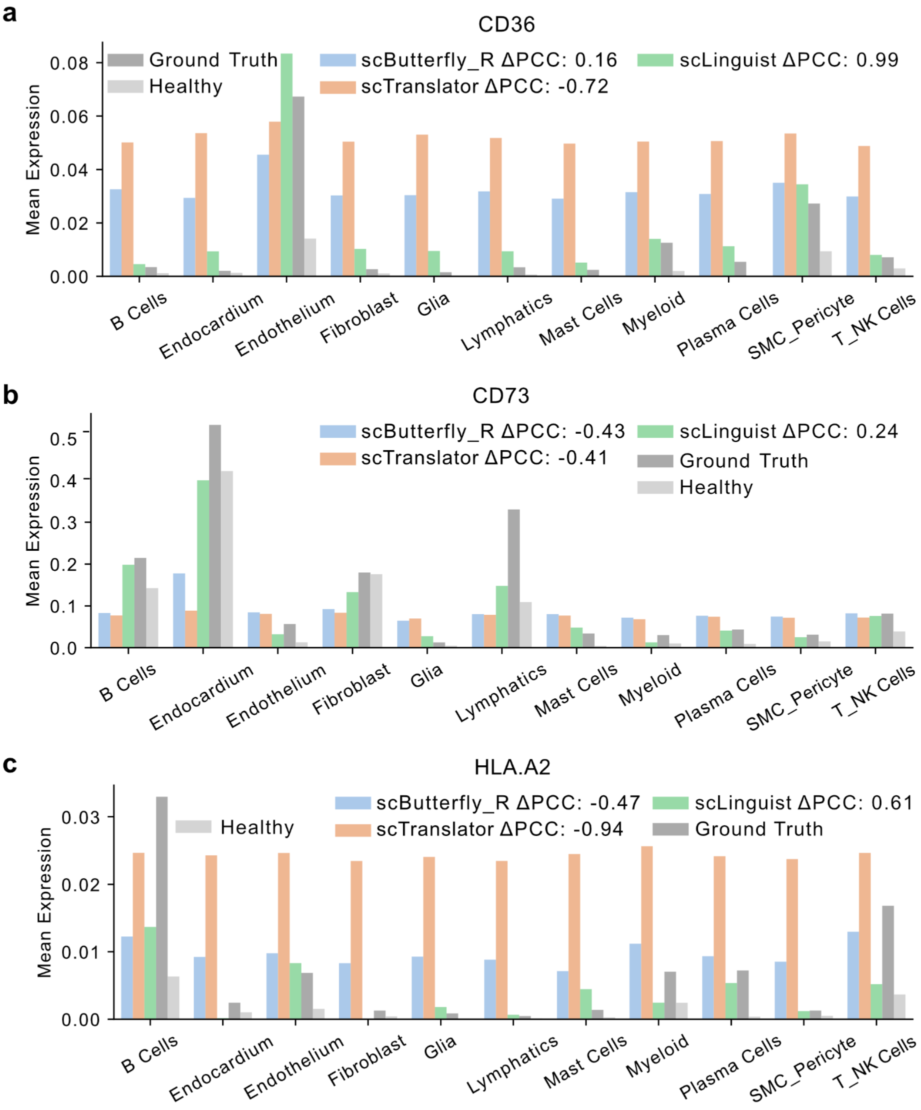


**Figure S10.** Comparison of predicted and true expression of disease-associated proteins across cell types. Bar plots comparing average protein expression across cell types for CD36, CD73, and HLA-A2, which are known to exhibit differential expression between healthy and heart failure conditions. scLinguist accurately captures the expression shifts, whereas other methods show less consistent trends.

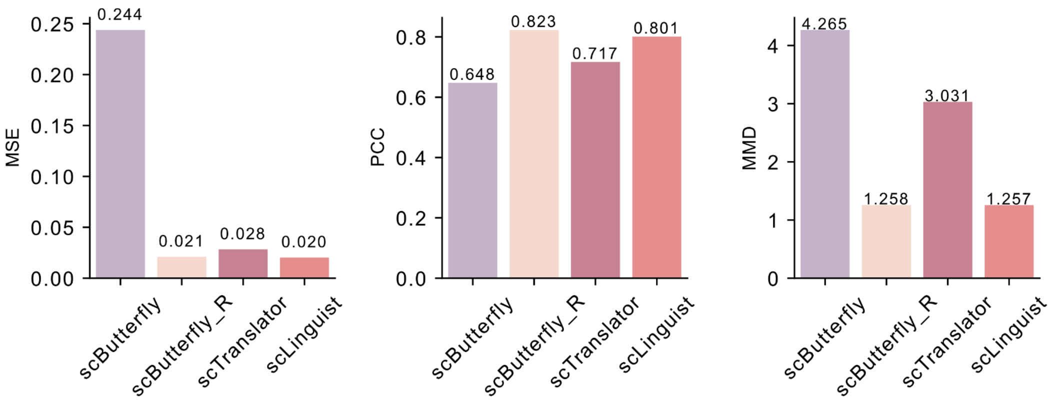


**Figure S11.** Evaluation of protein prediction on spatial transcriptomics data. Bar plots comparing scLinguist, scButterfly_R, scButterfly and scTranslator on Maximum Mean Discrepancy (MMD), Mean Squared Error (MSE), and Pearson Correlation Coefficient (PCC).

**Table S1**. Quantitative comparison of model performance under Setting 2a.

| Datasets | Metrics | totalVI | scArches | scButterfly | scButterfly_R | scTranslator | scLinguist |
|---|---|---|---|---|---|---|---|
| BM | MSE | 0.0073 | 0.0089 | 0.0805 | 0.0079 | 0.0086 | **0.0064** |
| BM | PCC | 0.9053 | 0.9058 | 0.8718 | 0.9369 | 0.9297 | **0.9469** |
| BM | MMD | 0.0295 | 0.0307 | 1.2814 | 0.0191 | 0.0111 | **0.0101** |
| BMMC | MSE | 0.0085 | 0.0097 | 0.0803 | 0.0083 | 0.0088 | **0.0078** |
| BMMC | PCC | 0.8018 | 0.8237 | 0.7535 | 0.8498 | 0.8319 | **0.8585** |
| BMMC | MMD | 0.0975 | 0.1637 | 0.2527 | 0.0875 | 0.0758 | **0.0751** |
